# Physiologically-relevant light exposure and light behaviour in Switzerland and Malaysia

**DOI:** 10.1101/2025.01.07.631760

**Authors:** Anna M Biller, Johannes Zauner, Christian Cajochen, Marisa A Gerle, Vineetha Kalavally, Anas Mohamed, Lukas Rottländer, Ming-Yi Seah, Oliver Stefani, Manuel Spitschan

**Affiliations:** Technical University of Munich, TUM School of Medicine and Health, Department Health and Sport Sciences, Chronobiology & Health, Munich, Germany; Translational Sensory & Circadian Neuroscience, Max Planck Institute for Biological Cybernetics, Tübingen, Germany; Centre for Chronobiology, University Psychiatric Clinics Basel (UPK), Basel, Switzerland; Monash University Malaysia, Electrical and Computer Systems Engineering, Intelligent Lighting Laboratory, Malaysia; Lucerne School of Engineering and Architecture, Lucerne University of Applied Sciences and Arts, Horw, Switzerland; TUM Institute for Advanced Study (TUM-IAS), Technical University of Munich, Garching, Germany

**Keywords:** Light Exposure, Cultural Characteristics, Lighting, Data Collection, Photometry, Wearable Technology, Radiometry, Subjective Exposure Assessment, Objective Exposure Assessment, Light Interaction

## Abstract

Light synchronises the internal clock with the external light-dark cycle. Keeping this alignment benefits health and prevents diseases. Quantifying light exposure is, therefore, vital for effective prevention. Since light exposure depends on photoperiod, culture, and behaviour, we investigated objective light exposure and individual light-related behaviour in Switzerland and Malaysia. In this observational field study, participants (N=39) wore a calibrated melanopic light logger at chest level for 30 consecutive days. At baseline and study end, the Pittsburgh Sleep Quality Index was assessed, and every 3 to 4 days, the Light Exposure Behaviour Assessment (LEBA) was filled. Our pre-registered analyses reveal that participants in Switzerland experienced brighter days (+3.16 times the average mel EDI) and spent more time (x1.9 times the duration) in daylight levels per hour of daylight, had ∼1.5h later bright light exposure in the afternoon, and stayed over 1h longer in dim light conditions before bedtime. LEBA scores did not differ between Malaysia and Switzerland, and LEBA items were stable over time. LEBA items also correlated with objective light exposure variables in Switzerland but not Malaysia, with a medium effect size (range of absolute r=0.32-0.48). These results highlight cultural and geographical differences in light exposure. We showed that LEBA can be related to actual light exposure and is ecologically informative, but this varies by culture.

## Introduction

Light is the main entrainment stimulus (zeitgeber) for synchronising human circadian rhythms to the natural light-dark rhythm of daylight given by Earth’s rotation (e.g, Foster et al., 2020; Kuhlman et al., 2018; Roenneberg & Klerman, 2019; Roenneberg & Merrow, 2016). A predictable and strong zeitgeber is key for a functioning sleep-wake rhythm and for human physiology to best deal with daily demands and stressors (e.g., Dijk & Archer, 2009; Foster, 2020; Windred, Burns, Lane, et al., 2024; Windred, Burns, Rutter, et al., 2024). Since the invention of electrical light in the 19^th^ century, the daily pattern of light exposure has rapidly altered from this robust light-dark cycle characterised by bright days and dark nights to a less predictable one, resulting from a potentially incalculable combination of artificial lights in the built environment and behavioural interactions with natural light and electrical light sources (Biller, Balakrishnan, et al., 2024). Additionally, bright days become dimmer due to a switch from daylight to weaker electrical sources indoors, while dark nights are brightened by electrical light (artificial light at night; ALAN). Factors that drastically influence natural exposure to daylight on a societal level are, to a large degree, the level of industrialisation and access to electricity (Pilz et al., 2018), and consequently, working hours such as shift work (Boivin et al., 2022).

Disrupting the entrainment of endogenous circadian rhythms by shifted or disordered light exposure can have various serious health consequences, such as disrupted sleep, shift work disorder, metabolic disorders, cardiovascular diseases, mood disorders, impaired immune system, or certain cancers (e.g., Ansu Baidoo & Knutson, 2023; Fishbein et al., 2021; Lunn et al., 2017; Neves et al., 2022; Roenneberg & Merrow, 2016). Recognising these negative consequences has led to a recent consensus statement on the optimal light exposure levels to best support physiology, sleep, and wakefulness in healthy adults (Brown et al., 2022). This consensus statement was strengthened through a concerted call to apply these recommendations in policy-making, lighting design and preventative measures for public and occupational health (Kervezee et al., 2024), leading to evidence-based statements for use in public health contexts (Spitschan et al., 2025). Analysing light exposure is, therefore, the necessary basis for restoring healthy, strong, and reliable circadian rhythmicity to both mitigate the negatives and increase the benefits.

However, the current expert recommendations by Brown and colleagues, albeit a beneficial and important start, especially for public health, are based on isolated laboratory lighting conditions and do not factor in individual light *behaviour* (Biller, Balakrishnan, et al., 2024). Photic history (Chang et al., 2013; Hébert et al., 2002; Smith et al., 2004) and an individual’s spectral diet (Webler et al., 2019) – the timing and spectral composition of prior light exposure – also play a role in subsequent sensitivity to light. Recommendations for healthier light exposure thus also need to factor in individual light exposure during day and night and how they interact with light sources. At present, there is minimal data about real-world light exposure and how much humans deviate from these target light levels. Previous studies often only sampled for a limited amount of time (e.g., several days to one or two weeks) and used wrist-based light loggers from actimeters (e.g., Didikoglu et al., 2023) or devices worn at spectacle frames (Stefani et al., 2024), the latter being valuable but limited in usability for more extended wear periods. Novel light logging devices at physiologically more relevant wearing positions (e.g. chest, eye level) are now available, thus quantifying ecological light exposure over long periods. One such device is the SPECCY light logger (Mohamed et al., 2021), a chest-worn wearable light logger that outputs numerous visual and non-visual metrics of light exposure following relevant standards from the CIE (International Commission on Illumination) standards via a spectral reconstruction algorithm (CIE, 2018).

Since individual light exposure-related behaviours also depend on an individual’s location (e.g., photoperiod, climate, temperature), culture, and the built environment, quantifying light exposure should factor in such behavioural and structural components (Biller, Balakrishnan, et al., 2024). The Light Exposure Behaviour Assessment (LEBA) instrument (Siraji et al., 2023) has recently been developed to quantify individual light-related behaviours. It categorises light exposure-related behaviours into five broad categories (factors) based on the participant’s recall of the past four weeks: i) wearing blue light filters, ii) spending time outdoors, iii) using phones and smartwatches in bed, iv) using light before bedtime, and v) using light in the morning and during daytime. This questionnaire is well suited to map objectively measured light exposure patterns to individual self-reported behaviours but also lends itself to studying potential cultural differences in interactions with light.

This project had three objectives. The first objective was to investigate differences in objectively measured physiologically-relevant light exposure light loggers at two sites that vary in culture and climate – Malaysia compared to Switzerland. A second objective was to investigate differences between the sites in subjectively measured light exposure behaviour using the LEBA questionnaire. The third objective was to examine which objectively derived light variables correlate with LEBA-derived items and composite scores and whether this depended on the study location.

Knowing more about the scope of assessment methods and cultural differences in light exposure and interactions with light can help inform future tailored and public health interventions to improve and foster healthy light exposure (Biller, Balakrishnan, et al., 2024).

## Methods

The current study was preregistered (Biller, Zauner, et al., 2024). We describe deviations from the preregistration document in the “Deviation from the preregistration” section below.

### Research questions and hypotheses

We addressed the following research questions and hypotheses as specified in the pre-registration document:

1. *Are there differences in objectively measured light exposure between Switzerland and Malaysia, and if so, in which light metrics?*

- **H1:** There are differences in light logger-derived light exposure intensity levels and intensity duration between Malaysia and Switzerland.
- **H0_1_:** No differences between Malaysia and Switzerland.
- **H2:** There are differences in light logger-derived timing of light exposure between Malaysia and Switzerland.
- **H0_2_:** No differences between Malaysia and Switzerland.
2. *Are there differences in self-reported light exposure patterns using LEBA across time or between the two sites, and if so, in which questions/scores?*

- **H3:** There are differences in LEBA items and factors between Malaysia and Switzerland.
- **H0_3_:** No differences between Malaysia and Switzerland.
- **H4:** LEBA scores vary over time within participants.
- **H0_4_:** No differences in LEBA scores over time among participants
3. *In general, how are light exposure and LEBA related, and are there differences in this relationship between the two sites?*

- **H5:** LEBA items correlate with preselected light-logger-derived light exposure variables.
- **H0_5_:** No correlation.
- **H6:** There is a difference between Malaysia and Switzerland in how well light-logger-derived light exposure variables correlate with subjective LEBA items.
- **H0_6_:** No differences between Malaysia and Switzerland.

### Study design

This observational study took place in Kuala Lumpur, Malaysia, and Basel, Switzerland, and was conducted for one month per participant at each site. Participants were not randomly assigned to treatment nor blinded in any way and were allowed to live their regular lives as usual. Day 0 and day 31 involved the collection and return of equipment, while days 1 – 30 involved the collection of light exposure and questionnaire data from the participants (**Fig. 1A**). Light data were collected via a neck-worn portable light logger (SPECCY). Questionnaires were deployed online via Qualtrics (Malaysia) and REDCap (Switzerland) (Harris et al., 2009, 2019)

**Figure 1.**
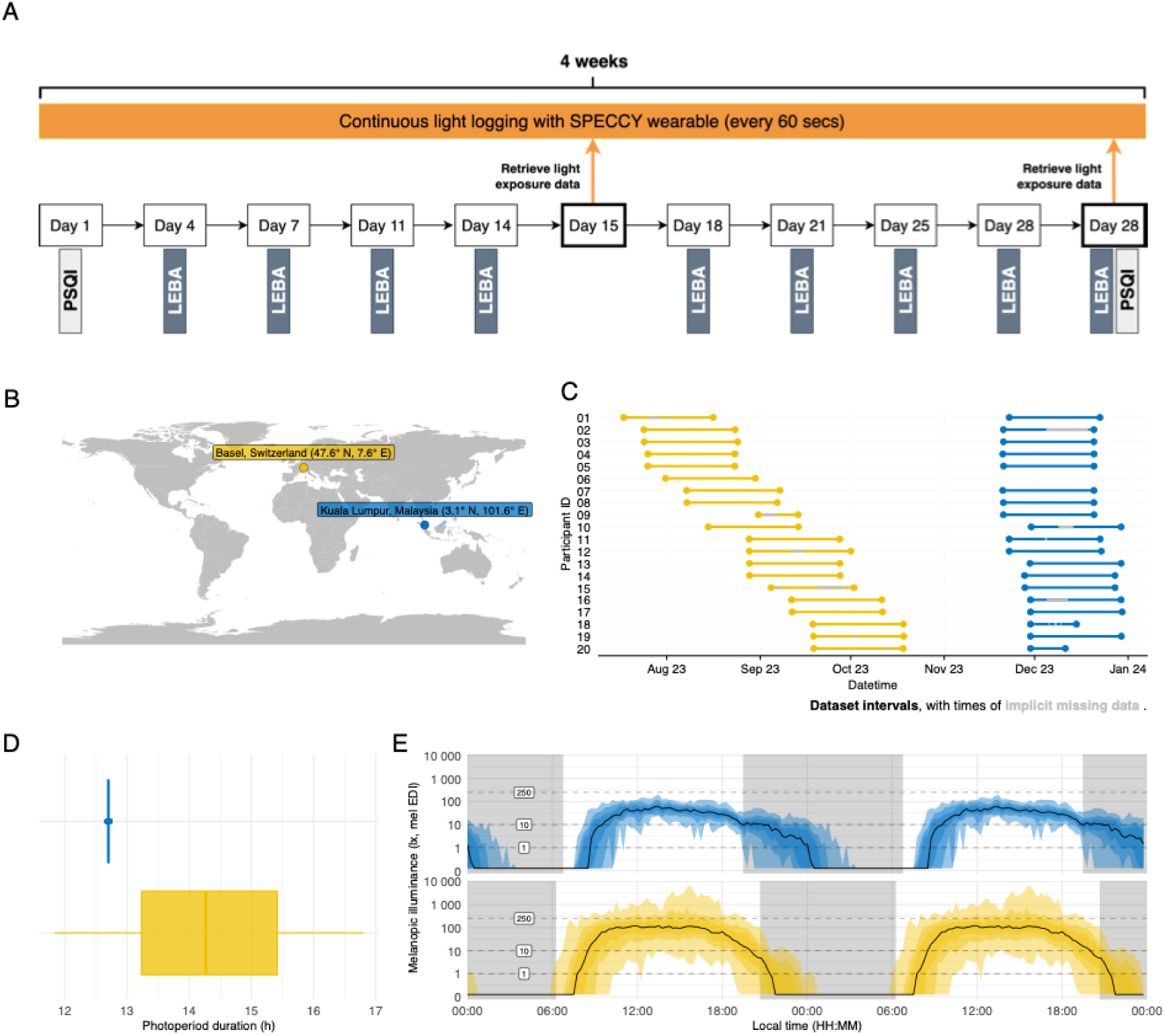
Overview of the study design. **A,** Shows the study design and measurements included for both research sites. **B**, Both research locations (Basel, Switzerland; Kuala Lumpur, Malaysia) shown on a world map. **C,** Individual recording periods per site and participant including implicit missing data in grey. Note that participant ID 06 is missing in the Malaysian dataset due to a lost light logger. **D,** Photoperiod duration in Malaysia (blue), and Switzerland (yellow). **E,** Grand averages of light exposure of illuminance (melanopic EDI) for Singapore (blue) and Switzerland (yellow). Abbreviations: LEBA, Light Exposure Behaviour Assessment; mel EDI, melanopic equivalent daylight illuminance; PSQI, Pittsburgh Sleep Quality Index.

On day 0, participants collected the light logger and filled in the Pittsburgh Sleep Quality Index (PSQI) questionnaire. The light logger was worn around the neck at chest level, approximately 20 cm below the wearer’s chin, with the sensing area facing outwards towards the sight of light arriving on the wearer’s eyes. From day 1 to day 14, participants wore the light logger continuously except when exposed to high amounts of water (e.g., bath, swimming) or when the dosimeter was being charged. Participants were asked to charge the sensor daily for an hour. The participants chose this charging time, but it was recommended to the participants that it remained consistent throughout the experiment and preferably at a time when they were primarily sedentary. Participants were also instructed to ensure that the sensor area of the light logger was not covered by any obstacles or objects (e.g., hands, hair, seatbelts, etc.). No specific instructions were given on light logger placement while participants were asleep. When the light logger was not worn, participants were instructed to place it so that its sensor area was blocked and could not detect any light. Participants were asked to answer the LEBA questionnaire twice a week before going to bed at night. On day 15, the light exposure data recorded by the light logger was downloaded from the device, and on day 31, participants returned the device, after which the collected light exposure data were downloaded. Data collection was completed after participants answered the Pittsburgh Sleep Quality Index (PSQI) questionnaire.

### Inclusion and exclusion criteria

In total, 20 (Switzerland) and 19 (Malaysia) participants were recruited to participate in the data collection experiment with the following self-reported (no standardised questionnaires) inclusion criteria:

1. Age between 19 and 65 years old
2. No history of drug abuse
3. Caffeine consumption ≤2 cups per day
4. Alcohol consumption <2 drinks per day or 14 drinks a week
5. Smoke <2 cigarettes per day

### Data collection procedures

Data collection took place at the Psychiatric University Clinics (UPK), Basel, Switzerland 47.5714° N, 7.5677° E) and at Monash University, Kuala Lumpur, Malaysia (3.0650° N, 101.6009° E), but participants were allowed to roam freely (**Fig. 1B**). Data collection in Switzerland was carried out between July and October 2023 (summer/autumn), while data collection in Malaysia was carried out between November and December 2023 (**Fig. 1C**). Photoperiod change during the collection period was considerable in Switzerland, but not Malaysia (**Fig. 1D**). Participants stayed within a 50 km radius of the respective sites.

For participants in Malaysia, applicants were asked to answer questions related to the inclusion/exclusion criteria using Google Forms. A total of 50 applicants were registered. Fourteen applicants did not meet the inclusion criteria, and the remaining applicants were either accepted, declined participation, or not contacted due to the maximal sample size being reached. One person was later excluded from the analysis due to a lost SPECCY device, leading to a final sample size of 19 participants in the Malaysian dataset. Qualtrics links to the LEBA and PSQI questionnaires were sent to the participants according to their schedules, and they would fill in the questionnaires via the link.

Participants in Switzerland were recruited through an online advertisement posted on the University’s bulletin board. The advertisement included a brief description of the study and inclusion and exclusion criteria. Those who expressed interest in the study were contacted via email to verify their eligibility. Upon confirmation, they were provided with a detailed study information sheet. Eligible and interested applicants were scheduled to visit the lab on a Monday. Participants were briefed in person about the study and registered using Qualtrics. Participants also completed the PSQI on-site using their mobile phones and received instructions on how to use the light logger.

### Wearable light logger

A wearable light logger, SPECCY, was used for data collection in Malaysia and Switzerland. The light logger, modified from the light spectral sensor developed by Mohamed et al. (2021) and developed at Monash University Malaysia’s Intelligent Lighting Laboratory (MMILL), captures spectral information across the visible range between 380 and 780 nm. The Australian Photometry and Radiometry Laboratory validated SPECCY, which has an effective measuring range from 1 lux to 130,000 lux. The light logger system is constructed from three multi-channel sensors, which provide 14 optical sensing channels within the measurement range of 380 nm and 780 nm and four channels in the infrared (IR) range. These channels capture low-resolution spectral data through proprietary software layers adjusted for sensor saturation and baseline signal calibration. Incorporated within a compact printed circuit board (PCB), the device houses the three multi-channel sensors, optimally placed for temperature consistency, an ambient temperature sensor, a microprocessor enabling Bluetooth Low Energy (BLE) connectivity, 16 MB of onboard flash storage allowing for over 130,000 spectral measurements, vital indicators and controls such as LEDs and a connectivity toggle button, and a micro-USB charging port. These components are securely encased within a 3D-printed housing made of polylactic material. A cosine corrector built into the case ensures appropriate sensor directionality. The output measurements are consistent with the CIE S 026/E:2018, including melanopic equivalent daylight illuminance (CIE, 2018).

### Surveys

#### LEBA

The Light Exposure Behaviour Assessment instrument (LEBA; Siraji et al., 2023) was developed to examine how light exposure behaviours affect health and well-being. LEBA categorises five key behaviours: wearing blue light filtering glasses indoors and outdoors (Factor 1), spending time outdoors (Factor 2), using phones and smartwatches in bed before sleep (Factor 3), controlling and using ambient light before bedtime (Factor 4), and using light in the morning and during daytime (Factor 5). **Table S1** lists detailed computations of these factors.

#### PSQI

The Pittsburgh Sleep Quality Index (PSQI; Buysse et al., 1989) is a self-report questionnaire that assesses sleep quality over one month. It measures seven components, including sleep duration, disturbances, and daytime dysfunction, and produces a global score. A higher global score (≥5) indicates poor sleep. **Table S2** lists details on the computation of components and global scores.

#### Sample size calculation

No specific sample size was calculated for the study. The sample was based on convenience sampling and resource limitations. No specific stopping rule was implemented.

### Measured variables

#### SPECCY-derived variables

Light data from the SPECCY device (melanopic EDI) were analysed using the R package *LightLogR v0.4.2* (Zauner et al., 2024) to perform cleaning steps, create visualisations, and calculate derived light exposure metrics. Melanopic EDI, the melanopic equivalent daylight illuminance (mel EDI) measured in lux, measures how strongly light affects the melanopsin system, which is the primary driver of the non-image-forming pathways in humans (Brown et al., 2022). An overview of each calculated light metric is given in **Table S3.** The exact calculation equation for each metric is contained within the open-source code of LightLogR (https://github.com/tscnlab/LightLogR).

#### LEBA-derived variables

The Light Exposure Behaviour Assessment instrument (LEBA; Siraji et al., 2023) captures light exposure-related behaviours on a 5-point Likert-type scale ranging from 1 to 5 (1 = never; 2 = rarely; 3 = sometimes; 4 = often; 5 = always). The score of each factor is calculated by the summation of scores of items belonging to the corresponding factor (**Table S1**). Respondents are requested to respond to each item retrospectively to capture their propensity for different light exposure-related behaviours. Originally, this covers the past 4 weeks but was amended in the current study to “past 3-4 days” (asked twice per week) for the Swiss site (the original framing of 4 weeks was kept for the Malaysian site). A list of all LEBA items is given in **Table S5**.

#### Composite scores

For the LEBA, five factors were calculated based on individual LEBA items (**see Table S1**). In scoring the PSQI, seven component scores are derived, each scored 0 (no difficulty) to 3 (severe difficulty). The component scores are summed to produce a global score (range 0 to 21). Higher scores indicate worse sleep quality, with a score >5 suggesting significant sleep difficulties.

### Transformations of variables and preprocessing steps

#### LEBA-derived variables

Item 4 of the LEBA questionnaire was reversed scored for the F2 calculation (see section “Measured variables” for more information on items and factors of the LEBA). LEBA scores were stored in a dummy-coded way in the Switzerland dataset (exported from RedCap), where 1 to 5 encode “Never”, “Rarely”, “Sometimes”, “Often”, and “Always”, respectively. Answers were directly coded in the Malaysia dataset (exported from Qualtrics). Upon import, all LEBA scores were converted to dummy-coded factors as described for the Switzerland dataset.

#### Sleep times and Chronotype from PSQI variables

Items 1 through 3 of the PSQI were used to calculate sleep/wake times and chronotype. Sleep onset was derived by adding typical bedtime (item 1) and sleep latency (item 2). For sleep offset, item 3 was used directly (wake time). For chronotype, the duration of sleep was calculated (sleep offset minus sleep onset on a circular scale), halved, and added to sleep onset. The resulting metric indicates the timing of mid-sleep as a proxy for chronotype (similar to chronotype, i.e. MSFsc, derived from the Munich Chronotype Questionnaire<u>; Roenneberg et al., 2019)</u>. As free and workday information were not collected, no sleep correction for sleep debt (as done to calculate MSFsc from the MCTQ) during working days was applied.

#### Melanopic EDI-derived variables

The base measurement was the time series of melanopic EDI (in lux). Several summary metrics are calculated from that measure (see **Table S3**). The period over which these metrics are calculated varies depending on the specific research question, hypothesis, and metric. Generally, metrics were calculated per participant and day, except interdaily stability (IS), which was calculated per participant. Whenever melanopic EDI was directly used in a statistical model, it was logarithmically transformed (base 10).

#### Time-of-Day

For some research questions, the distinction of daytime vs. evening or nighttime was relevant when calculating metrics, as either time of day was part of the statistical model or only a portion of the day was relevant. In those cases, metrics were calculated based on a filtered time series of melanopic EDI. These filters have three thresholds: dusk, dawn, and midnight. Dusk and dawn were calculated based on latitude, longitude (Switzerland: 47.5585° N, 7.5839° E; Malaysia: 3.0650° N, 101.6009° E), and calendar date with the *suntools* package (Bivand & Luque, 2023), with a solar depression angle of 6°, which yields civil dusk and dawn. Dawn to dusk is considered daytime, dusk to midnight as evening, and midnight to dawn as nighttime.

#### Datetimes

Since the two sites were in different time zones, directly including the datetime variable in a statistical model or a combined visualisation was not sensible since we were interested in differences depending on local time, not real-time differences due to different time zones. When only the local clock time (not the date) was of interest, datetime was transformed into seconds from midnight local time. When the local time, including date, was of interest, datetime was transformed by overwriting the time zone of both sites to UTC, thus forcing them to operate on the same timeline.

#### Correlating LEBA scores with light exposure metrics

Depending on the site, different timespans were used in the correlation analysis (**H5**) due to how the items in the questionnaire were framed:

For Switzerland, the LEBA covered behaviour in the current respective period (i.e., about the past three or four days), and light exposure metrics were calculated from the timespans between each LEBA questionnaire. These metrics were correlated with the respective LEBA scores. For example, LEBA scores from day 7 were correlated with light exposure metrics from days 5 to 7, as the prior LEBA questionnaire had been collected at the end of day 4. All other time spans between assessments were calculated accordingly.

For Malaysia, where the LEBA had asked about the behaviour in the past four weeks, light exposure metrics were calculated for the entire four weeks of data collection and correlated with the final round of LEBA scores collected on day 31. An average value was calculated prior to correlation for light exposure metrics that were calculated for each day. This avoided a one-to-many comparison (one LEBA datapoint to many light exposure metric datapoints) that would have skewed significance tests of the correlation.

### Descriptive statistics and statistical models

We also tested if light exposure and LEBA items were correlated (**RQ3**) by calculating specific light variables using the R package *LightLogR* and correlated them to the respective LEBA item. See **Table S5** for an overview of which light variables were calculated for each LEBA item.

#### Exploratory analyses

We extended our analyses beyond the specified hypotheses described above in a secondary analysis. Since we expected differences between the two sites regarding objectively measured light exposure (using the SPECCY light logger; **RQ1**) and subjectively measured light exposure (using the LEBA questionnaire; **RQ2**), we also wanted to test this thoroughly in a data-driven way. To this end, we did a cross-correlation of all available light metrics in *LightLogR* for the objectively measured light exposure with all available variables and factors from the LEBA questionnaire across sites. We further looked into dependencies of light exposure and derived metrics due to age, sex, and chronotype. Lastly, we assessed the compliance with the recommendations for healthy day, evening, and night light exposure (Brown et al., 2022).

#### Inference criteria

##### Check for assumptions

Appropriate model diagnostics were performed for all statistical tests to ensure linearity of (generalised) relationship, normally distributed residuals and random effects, and homoscedastic residuals. In the case of generalised additive models, the number of knots (k) was tested to ensure sufficient degrees of freedom to adequately capture the nonlinear relationships.

##### Statistical tests

**Table S4** links specific statistical tests to the research questions and hypotheses. Overall, one of five statistical model types was used:

1. (Generalised) linear mixed-effect models for continuous numeric dependent variables, using the lmer() and glmer() functions in the *lme4* package version 1.1-35.5 (Bates et al., 2015). Generally, a Gaussian error distribution was assumed for dependent variables. A Poisson error distribution was used for count data, such as the seconds above a given threshold.
2. Cumulative link mixed-effect models for ordinal dependent variables, using the clmm() function in the *ordinal* package version 2023.12-4.1 (Christensen, 2010).
3. Generalised additive mixed-effect models for non-linear relationships, using the gam() and bam() functions in the mgcv package version 1.9-1 (Wood, 2000)
4. Correlation matrices using Spearman-rank correlation for the correlation of continuous numeric and ordinal variables. These models use the cor() function in the *stats* package version 4.4.2 in base R (R Core Team, 2021) and the rcorr() function in the *Hmisc* package version 5.2-1 (Harrell, 2003). For the correlation of chronotype (mid-sleep) with the mean timing of light above threshold (MLiT250), we used circular correlation with the *circular* package, version 0.5-1 (Lund & Agostinelli, 2004).
5. Bootstrapping to determine 95% confidence intervals for variables. Bootstrapping was performed with base R functions such as sample() and the *tidyverse* package version 2.0.0 (Wickham, 2016; Wickham et al., 2019).

##### Accounting for multiple testing

A false-discovery-rate correction (Benjamini-Hochberg) was applied for every hypothesis based on the number of models/tests performed within that specific hypothesis.

##### Effect size

For the statistical tests in model types 1, 2, and 3, unstandardised effect sizes are reported as a result of the final models, i.e., beta-coefficients and non-linear dependency curves. For type 4, the correlation coefficients are effect sizes.

##### Significance levels

For statistical tests in model types 1, 2, and 4, p-values were calculated and corrected for multiple testing. Values equal to or below 0.05 were considered significant. For type 1 and type 2, p-values are calculated through a likelihood ratio test of models with and without a parameter in question. For type 3, model selection/significance was based on Akaike’s Information Criterion (AIC), where the more complex model was preferred if their AIC was two or more below the less complex model. For type 4, p-values were calculated through the asymptotic t-approximation (Press et al., 1992). For type 5, 95% confidence intervals were calculated based on 10^4^ bootstrapping samples. If a standard deviation of 0 was not within the confidence interval, the item variance over time was considered significant.

##### Photoperiod correction

**Fig. 1D** shows the differences in photoperiod between the two sites during data collection. The photoperiod, as measured from civil dawn to civil dusk, was, on average, 12.7 hours in Malaysia during data collection, while it was 14.4 hours in Switzerland. Furthermore, while there is negligible difference in photoperiod within the Malaysia site, there is a strong variation within the Switzerland site due to higher latitude and a longer data collection period (not per participant but for overall data collection). To account for this, metrics in H1 were corrected by their respective periods: times above or below a given threshold per day were divided by the (photo)period relevant for the metric. The resulting metrics should be interpreted such that per hour of photoperiod, *x* seconds were spent above/below the threshold. Similarly, this correction was applied for time metrics covering the evening, which were divided by the evening length.

### Data exclusion, outlier handling and awareness checks

#### Exclusion criteria of data

Measurement data were excluded if their values lied outside the measurement equipment’s plausible thresholds (≥130,000 lux for photopic illuminance). No awareness checks were implemented during data collection, and outlier handling was not necessary after viewing the data.

#### Missing data

The missing data percentages (implicit missing data) and percentage data where the recorded photopic illuminance fell outside the light logger’s effective measuring range between 1 lux and 130,000 lux (explicit missing data) were recorded and analysed. Values above the upper threshold were few (one data point for Malaysia and eight data points for Switzerland) and negligible relative to the amount of available data. Values below 1 lux were regular and expected, as this value indicates darkness. Values below 1 lux were converted to 0.1 lux, which allowed for their inclusion in logarithmically transformed models.

All implicit missing data were made explicit in the analysis, i.e., gaps in the time series were filled with the respective date, and corresponding measurement values were set to NA. Participant days with more than a fixed threshold of 6 hours of missing data per day were excluded from further analysis. The threshold was determined by a sensitivity analysis using three randomly chosen light exposure metrics and three randomly selected participants without missing data. The threshold was set so that the average of each metric stayed within a 95% confidence interval based on their original variation (mean±2 SE) and 10^4^ resamples for each threshold. This ensured a threshold that did not significantly affect metric calculation based on the actual data (see **Fig. S1**). While the threshold may seem rather large, it exemplifies that for a sufficiently long time series, i.e. 30 days in this case, the chosen light exposure metrics (IV, last timing below 10 lux, MP ratio) are rather robust to missing data, even for larger time frames within a day. For an overview of missing data, see **Table S6**.

### Deviations from the preregistration

In research question 1, H2 *(“There are differences in light logger-derived timing of light exposure between Malaysia and Switzerland.”*) of the preregistration document specified an additive model to describe light exposure patterns within participants and between sites. This approach led to the conclusion that the light exposure patterns were dominantly driven by individual patterns rather than an overarching pattern by site (see **Results**). To explore differences by site, another model that only included random intercepts by participant (rather than random smooths by participant) was constructed. Both models are part of the analysis documentation in the **Supplementary Materials (Section 5.2.2.4),** and their results will be discussed according to their respective relevance.

The preregistration document specified a generalised mixed model approach to test H6 (“*There are differences between Malaysia and Switzerland on how well light-logger derived light exposure variables correlate with subjective LEBA items”*). As the specified formula did not lead to the stated number of models, i.e. 23+5, and also did not provide an overall overview of whether selected light exposure metrics correlate with LEBA items and factors, we provide here an alternative approach by analysing whether there are significant differences in correlation coefficients between the sites. In this approach, a linear model is used, and the number of models equals the number specified in the preregistration. The output is also more suitable for answering the hypothesis. The stricter analysis is still included in section 5.4.2. of the analysis documentation (link in the supplementary material).

We also had to correct H0_4,_ which originally stated “*No differences between Malaysia and Switzerland*”. However, this comparison is not sensible given that we were investigating the LEBA scores within the Malaysian dataset over time instead of investigating if these scores differed over time, dependent on the research site. The updated H0_4_ is thus: “*No differences in LEBA scores over time within participants*”, as already stated above at the beginning of the Methods section.

## Results

### Participant characteristics

The sample in Basel, Switzerland, consisted of five male and 15 female participants (age 30.35 ± 9.7 years old), and all participants completed their data collection successfully. In Malaysia, 8 males and 11 females (age 24 ± 6.7 years old) completed data collection successfully (**Fig. S3**), and one participant failed to complete the experiment due to the loss of the light logger. Age differed significantly between sites *(p = 0.03*), but there was no significant dependency of age on sex (*p = 0.97*). Sex did not differ significantly by site (*p = 0.32*), nor was there a significant difference in the number of male vs. female participants overall (*p = 0.98*). PSQI scores were comparable between sites, both on day 1 and day 31, and did not differ significantly (**Table S13-14**).

Timing of mid-sleep derived from the PSQI (as a proxy for chronotype) was 4:57 a.m. ±1.4 hours in Malaysia, and 3:45 a.m. ±0.8 hours in Switzerland. The difference is significant (*p = 0.003*; see **Fig. S5**). This difference of 1.2 hours in mid-sleep compares to a 0.37 hour difference in noon timing (i.e., daytime centre).

### Individual differences in light exposure

**Fig. 2**. shows individual light profiles from both sites averaged across the recording period. While the results below go into detail about condensed quantifiers of light exposure metrics, individual patterns lead to valuable insights into the wide breadth of light exposure patterns and their variation. To highlight just a few, some participants, such as ID 16 or 18 in the Swiss dataset, show very *small deviations* from their daily light exposure patterns as indicated by narrow 95-, 75-, and 50-percentile ribbons. On the other hand, participants 15 and 17 in the Swiss dataset or participants 08, 13 or 16 in the Malaysian dataset show *strong deviations* from their median daily patterns. Interestingly, this is somewhat captured by the respective interdaily stability (IS) measures but not as much as expected. In general, the IS values are also rather small (all <0.3), indicating strong variation between days (higher numbers mean more stability).

**Figure 2.**
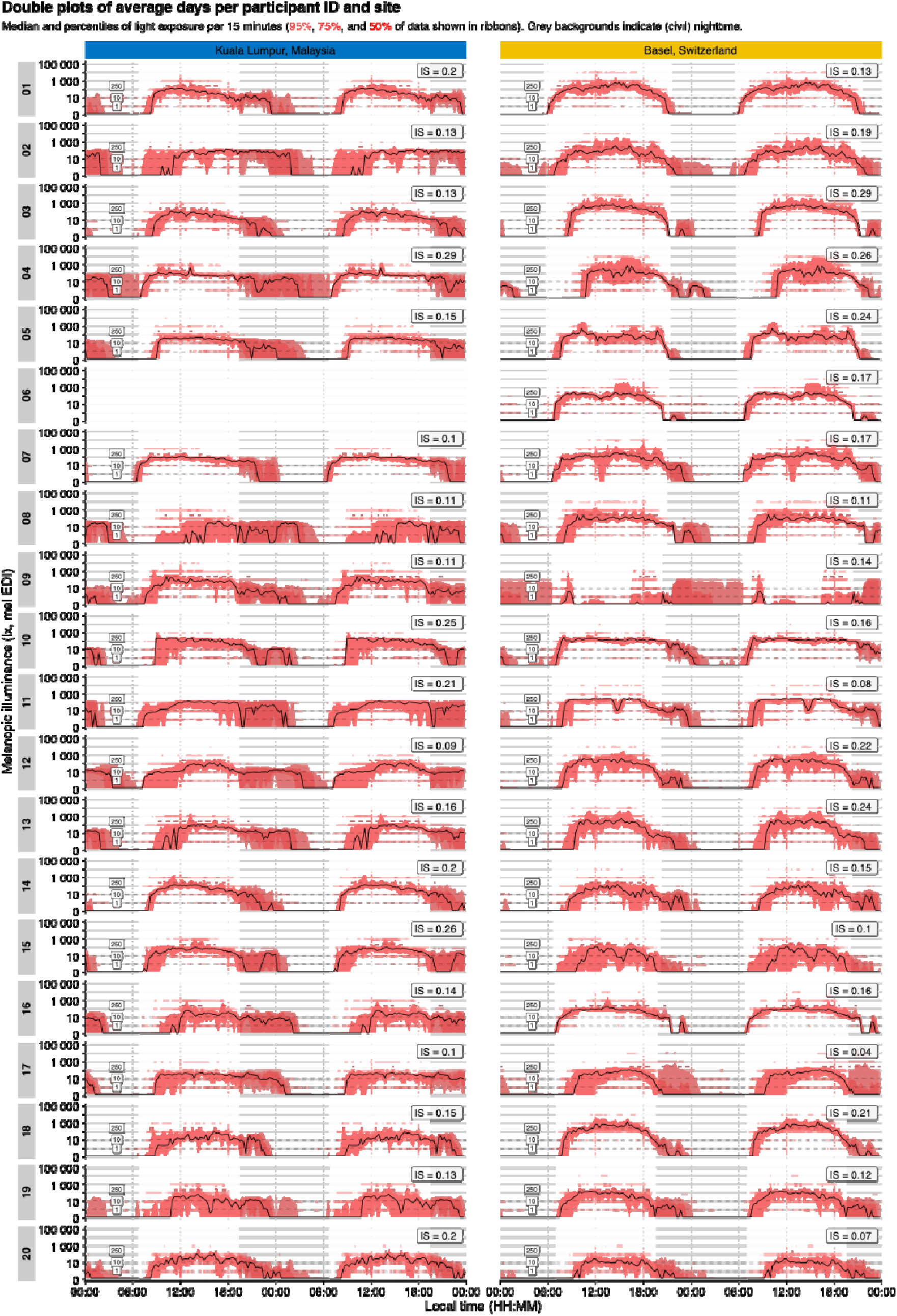
Double plots of average light exposure over participants across both research sites. Median and percentiles of light exposure measured in illuminance (lux, melanopic EDI per 15 minutes (95%, 75%, and 50% of data shown in ribbons). Grey background indicates civil nighttime. Abbreviations: IS, Interdaily stability. Melanopic EDI, Melanopic equivalent daylight illuminance.

Furthermore, the data also show a degree of within-day variation: while participant 08 in Malaysia shows a clear tendency for predominant nighttime exposure, there are regular “daytime patterns” visible in the 95- and 75-percentile ribbons. Participant 17 in Switzerland, on the other hand, has a clear daytime exposure profile but with regular evening and nighttime extensions and variations.

### Participants in Switzerland spent more time in daylight, experienced brighter days and stayed longer in recommended light levels during the day compared to participants in Malaysia

**Fig. 1E** shows the grand average of light exposure profiles for the site in Switzerland (yellow line) and the site in Malaysia (blue line). The model summary (**Table S7**) shows that participants in Switzerland spent significantly more time above threshold per hour of daylight for 250 lux (TAT250) and 1000 lux (TAT1000) than participants in Malaysia, i.e., x1.78 and x1.94, respectively (x1.77 over 250 lux melanopic EDI during daytime hours, TATd250). The second model summary (**Table S9**) also shows that the 10 brightest hours (M10m) of participants in Switzerland are significantly brighter than for participants in Malaysia, i.e., 661 lux and 229 lux, respectively. The frequency of crossing 250 lux is about twice as high for participants in Switzerland compared to Malaysia (64 absolute times compared to 36 times, respectively). Lastly, participants in Switzerland experienced light levels of above 250 lux melanopic EDI about 1.5 hours longer across the day, compared to Malaysia (19:09 vs 17:41 respectively, **Table S9**) and about 2.5 times higher melanopic EDI illuminance levels daily (437 lux vs 170 lux respectively) as demonstrated in the interaction of site and time of day (**Table S8**).

### Participants in Switzerland experienced earlier dim light exposure and darker evenings

Participants’ last time above 10 lux after dusk (LLiT10) in Switzerland was about 1 hour earlier than participants in Malaysia (22:05 vs 23:16, respectively). There was no difference (p<.9) in time below threshold of 10 lux during the evening (TBTe10) between sites (**Table S7**). Exploring the interaction of site and time of day (day vs evening) in illumination levels revealed an average for participants in Switzerland of 3.5 lux during evening hours and a 5-fold increase (17.6 lux) for participants in Malaysia (**Table S8**).

### Participants in Switzerland followed recommendations for healthy light exposure about half of the time, in Malaysia about 40% of the time

Using PSQI-derived typical sleep and wake times to assess whether participants followed the Brown et al. (2022) recommendations for healthy light exposure reveal are marked difference between both the categories and sites, which align well, however, with the results above regarding daytime and evening comparisons. The lowest level of compliance to the recommendations was for daytime levels (wake until three hours prior to sleep), where participants in Malaysia only managed to reach 250 lx melanopic EDI 10% of the time, compared to 26% in Switzerland. This increased markedly in the evening (last three hours before sleep), where Malaysian participants stayed at or below 10 lx melanopic EDI 64% of the time, compared to 78% in Switzerland. Better still were nights (sleep hours), with 84% of Malaysian participants at or below 1 lx melanopic EDI, compared to 92% in Switzerland. Overall, participants in Malaysia followed the recommendations 39% of the time, compared to 54% in Switzerland (see **Table 1**). A more granular depiction of when participants followed the recommendations can be found in **Fig. S6**.

**Table 1.** Compliance to Brown recommendations for healthy light exposure. Data covers all valid measurement periods for all participants. Daily values for *day*, *evening*, and *night* were averaged per participant and then per site, i.e., these represent an average participant-*day/evening/night* per site. *Overall* values represent time-weighted averages of *day*, *evening*, and *night*. I.e., values represent an average (full) participant-day per site.

| Compliance to Brown recommendation for healthy light exposure |  |  |  |  |
| --- | --- | --- | --- | --- |
|  | day | evening | night | overall |
| Malaysia | 10% | 64% | 84% | 39% |
| Switzerland | 26% | 78% | 92% | 54% |

*day* includes the time from wake until evening (recommendation $\geq 250\text{lx mel EDI}$ ). *evening* starts three hours prior to sleep until sleep onset (recommendation $\leq 10\text{lx mel EDI}$ ). *night* covers the sleep times (recommendation $\leq 1\text{lx mel EDI}$ ). *overall* indicates the time-weighted average compliance.

### Chronotype relates to mean timing of light in Malaysia, but not Switzerland

We found a slight correlation of mean timing of light above 250 lx (MLiT_250_) with chronotype (mid-sleep) in Malaysia (r = 0.22, *p <0.001*), but not in Switzerland (r = 0.05, *p = 0.27*; see **Fig. 3**). In a mixed-model framework, the overall dependency of MLiT_250_ on chronotype is significant (*p = 0.01*), but the site-specific interaction is not supported (*p = 0.35*). Thus, there is an overall prediction of 0.22 hours increase in MLiT_250_ per hour increase in mid-sleep.

**Figure 3.**
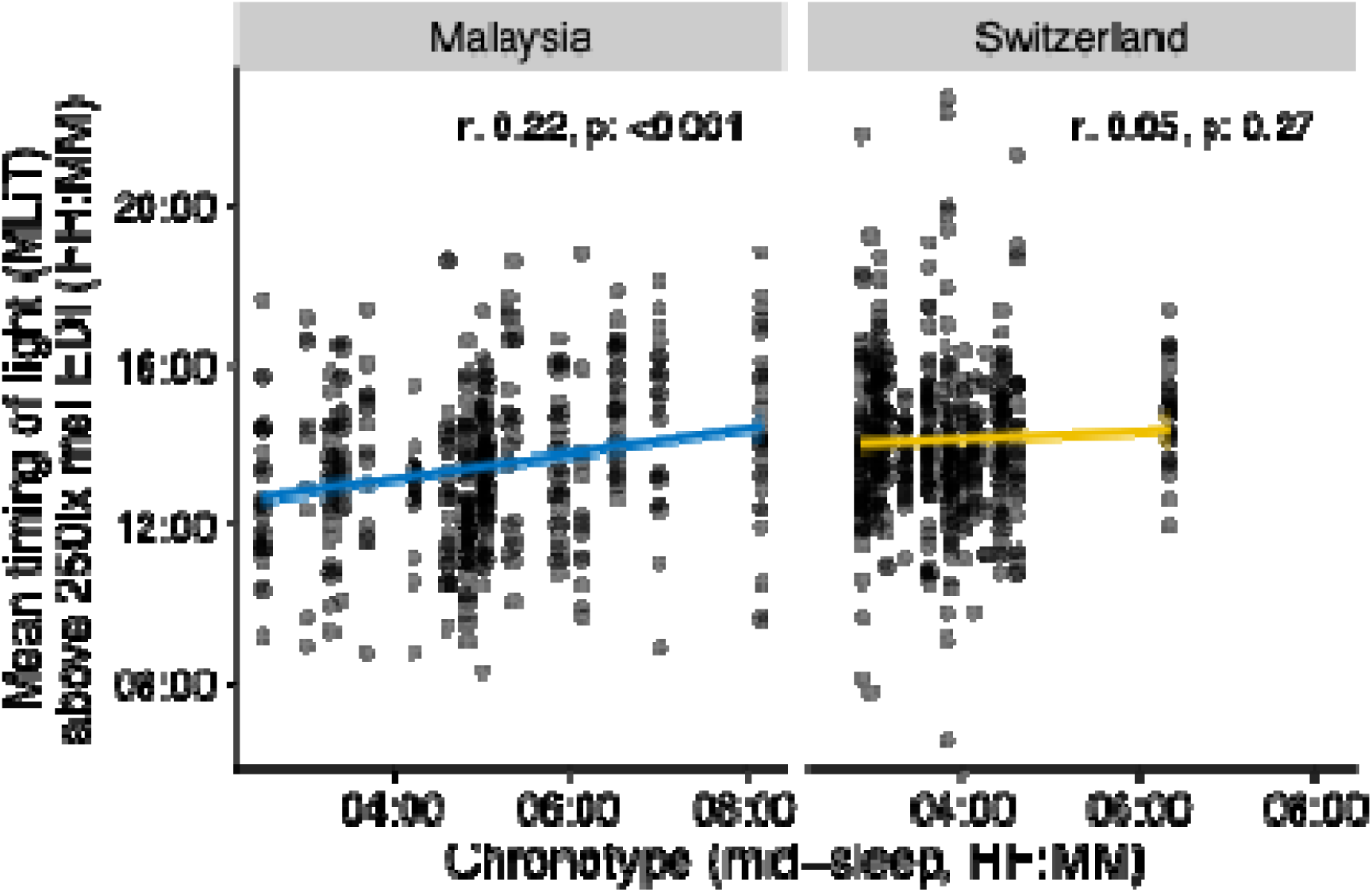
Chronotype influence on light exposure timing. Dependency of the mean timing of light above 250 lx melanopic EDI (MLiT_250_) against chronotype (mid-sleep) for each site. Dots indicate value-pairs, the line and bands a linear model of the dependeny. Correlation values (r-values) at the top of each panel show the (circular) correlation coefficient of the dependency, including its p-value.

### Modelled light exposure patterns vary between sites, with strong differences between individuals

**Fig. 4** shows model results when estimating light exposure patterns, with **Fig. 4A** showing individual patterns, **Fig. 4B** patterns by site, and **Fig. 4C** differences between the sites. While each site shows a characteristic pattern (**Fig. 4B**), these are composed of strongly varied individual patterns within each site (**Fig. 4A)**. In fact, when modelling each participant’s pattern, an overall site-specific pattern does not significantly increase the model’s explanatory power. This is indicative of strong differences between participants within each site. Overall, the light exposure patterns are consistent with the results derived from individual light metrics: Participants in Switzerland experience longer and brighter days and darker and earlier nights. While the longer photoperiod in Switzerland might partly explain the daytime effects, it does explain the described night-time effects.

**Figure 4.**
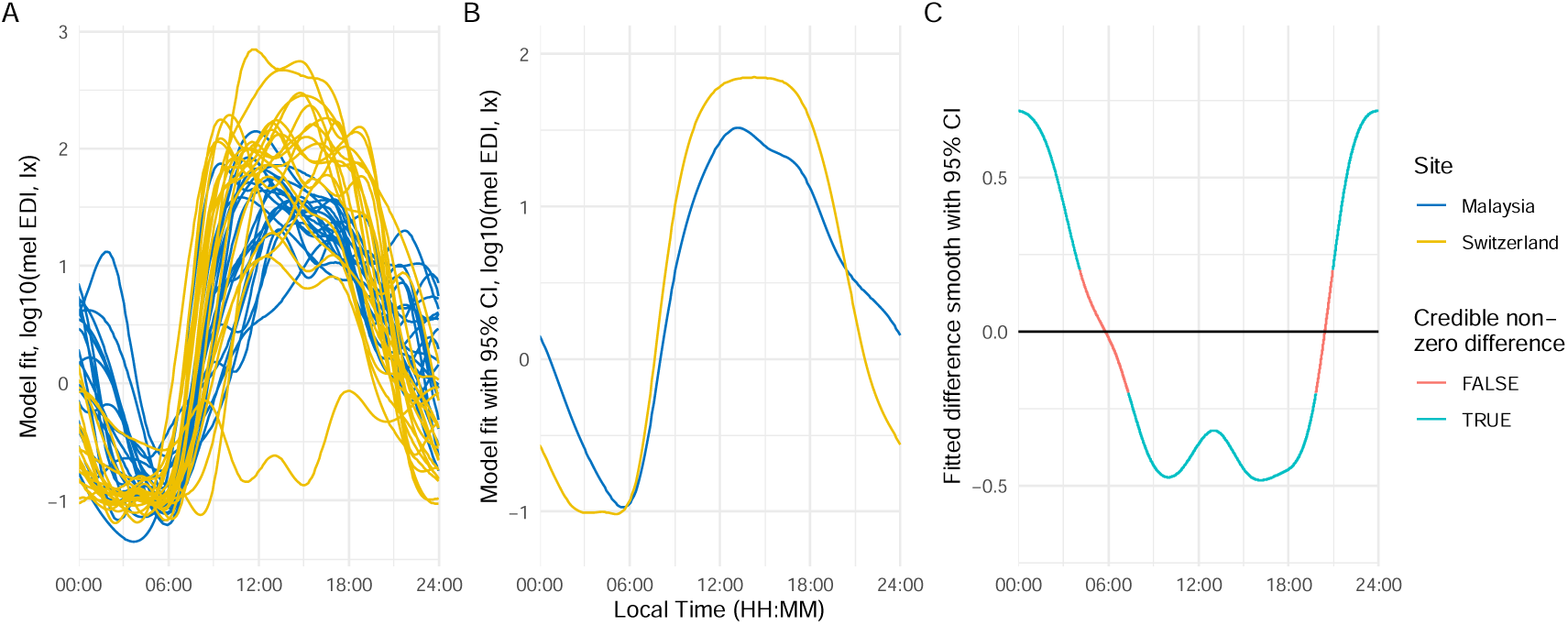
Model fits of light exposure differences between Malaysia and Switzerland. **A,** Individual model fits per ID and research site for melanopic EDI (lux). **B,** Model fits over all IDs per research site with with 95% confidence interval. **C,** Fitted differences between research sites smoothed with 95% confidence interval with an indication of significant differences per time. Abbreviations: CI, confidence interval; mel EDI, melanopic equivalent daylight illuminance.

Despite this strong individual variation, we can still reject H0_1_ and H0_2_ of our first research question since we did find statistical differences in light-logger-derived light exposure intensity levels and duration of intensity, as well as timing of light exposure, between Malaysia and Switzerland (see also **Table S10**).

Furthermore, there is strong evidence for a site-specific effect of sex. While there is no difference in overall melanopic EDI due to sex, the patterns of exposure differ. Specifically, females in Malaysia are less exposed to light in the hours before noon and around midnight, compared to males. In Switzerland, females are exposed to more light before noon, compared to males, and are also more likely to have a higher exposure in the afternoon. The effect size is a reduction of about 60% for women in Malaysia in the specified times of day, and about a 5-fold increase for women in Switzerland before noon, and about 2- to 3-fold in the afternoon. See **Fig. S2** for further details on sex-dependent patterns and differences.

Because of the significant age difference between sites, age is a confounder for a comparison between the sites (see **Fig. S3**). Thus, we did not explore how age relates to light exposure in general, but rather whether age differences from the site-mean affect light exposure. These results will have to be taken very tentatively, as the data is sparse. Even when corrected for the age difference, the overall difference pattern between the sites remained: Participants in Switzerland exhibit brighter days, particularly in the morning, and darker nights. There is some evidence that younger participants (5 years below average) in Malaysia have brighter mornings and darker nights, whereas younger participants in Switzerland have brighter times around and after noon, as well as darker evenings, each compared to their average age group. Results for older participants (20 years above site average) is not reported here, as there are only one and three participants in that age group in Malaysia and Switzerland, respectively, prohibiting generalisations. Results are still shown in the full analysis document (https://github.com/tscnlab/BillerEtAl_RSocOpenSci_2025).

Finally, light exposure patterns differ due to photoperiod, which could only be analysed in the Swiss dataset (see **Fig. 1)**. Unsurprisingly, light exposure is higher and ends later in the day when photoperiods are longer. However, while the shift in exposure end is somewhat pronounced, it is very moderate in the morning. Across all photoperiod durations, there are two high points during the day, one around noon and the second in the afternoon around 4 p.m., with dips in between (see **Fig. S4**).

### LEBA responses do not differ between Malaysia and Switzerland and are stable over time

To address the second research question, we compared LEBA items and scores across sites and found that under the strict p-value adjustment for H3 (n=28), none of the five LEBA factors or individual items were significantly different between sites (for all: p >.3; see section 5.3.1 in the full analysis document linked in supplementary material). Thus, we cannot reject our H0_3_ for this research question since we could not find strong differences in Malaysian and Swiss LEBA scores.

Looking at the stability of LEBA scores across time within the Malaysia site (H4), bootstrapping analyses showed that scores were very stable over one month. All 23 questions and 1 out of 5 factors did not vary significantly in more than 50% of participants (Table S11). We consequently cannot reject H0_4_ and assume scores to be stable over time, at least in this dataset (see Table “Average change in LEBA metrics across four weeks of data collection” in section 5.3.2. of the full analysis document linked in the supplementary material).

### LEBA items correlate with preselected light-logger-derived light exposure variables in Switzerland but not in Malaysia

While exploratory analysis shows correlations in both sites, only Switzerland shows significant correlations after correction for multiple testing (**Fig. 5**). Out of 84 correlations hypothesised *a priori*, 10 were significant in Switzerland.

**Figure 5.**
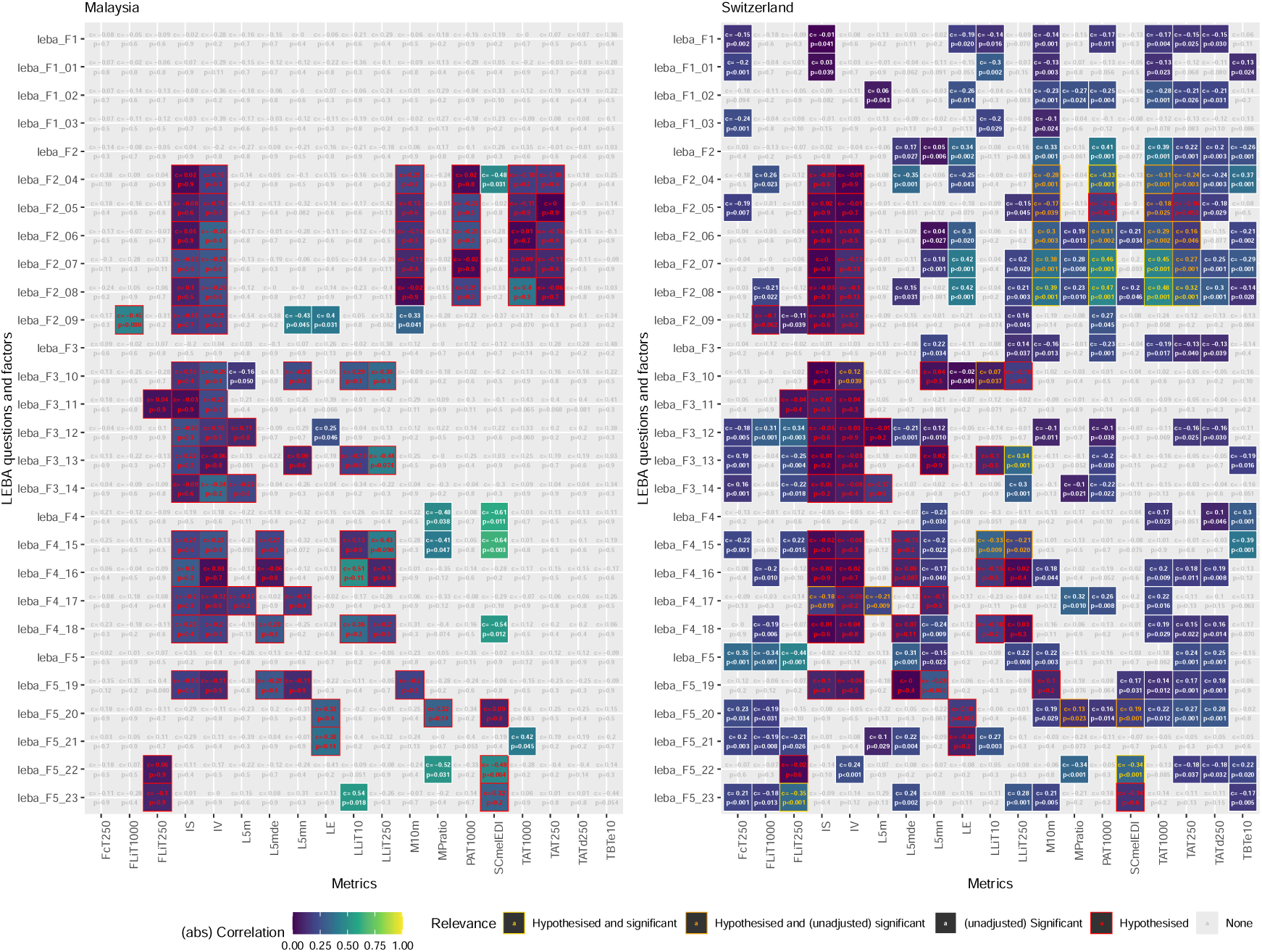
Correlations between individual LEBA items and pre-selected light metrics for Malaysia and Switzerland. Abbreviations: FcT250, Frequency crossing threshold at 250 lux; FLiT1000, Frequency of light exposure above 1000 lux; FLiT250, Frequency of light exposure above 250 lux; IS, Interdaily Stability; IV, Intradaily Variability; L5m, Mean light exposure during the least active 5 hours; L5mde, Mean duration of exposure during the least active 5 hours; L5mn, Number of exposures during the least active 5 hours; LE, Light Exposure; LEBA, Light Exposure Behaviour Assessment; LLiT10, Lowest light exposure intensity threshold at 10 lux; M10m, Mean light exposure during the most active 10 hours; MPratio, Ratio of mean photopic to melanopic light exposure; PAT1000, Period above threshold at 1000 lux; SCmelEDI, Spectral composition of melanopic equivalent daylight illuminance; TAT1000, Time above threshold at 1000 lux; TAT250, Time above threshold at 250 lux; TATd250, Time above dynamic threshold at 250 lux; TBTe10, Time below threshold at 10 lux.

The effect size of those correlations was medium on average (r = 0.39), and all correlations showed the hypothesised direction (**Table 2**). We can thus reject H0_5_ since we did indeed observe statistically significant correlations between LEBA and light-logger-derived variables There were also a few LEBA questions where the correlation coefficients differed significantly between sites. Specifically, these were the questions *“I dim my mobile phone screen within 1 hour before attempting to fall asleep”* (F4 item 15) and *“I dim my computer screen within 1 hour before attempting to fall asleep”* (F4 item 18; **Fig. 5**). While the correlation is positive with preselected light exposure metrics in Malaysia (r= .25 both, p=.019), it is zero or negative in Switzerland (r = -.01 and -.15, respectively, p=.035; **Table S12**). We can, therefore, also reject H0_6,_ given that we observed differences in the strength and directions of LEBA scores/items and light-logger-derived variable correlations between Malaysia and Switzerland.

**Table 2.** Significant correlations between individual LEBA items and hypothesised light metrics for Malaysia and Switzerland. P-values are considered significant below 0.05. Abbreviation: FLiT250, First time above threshold of 250 lux; LLiT250, Last time above threshold of 250 lux; M10m, Mean light exposure during the most active 10 hours; PAT1000, Period above threshold at 1000 lux; SCmelEDI, Spectral composition of equivalent daylight illuminance; TAT1000, Time above threshold at 1000 lux; TAT250, Time above threshold at 250 lux.

| Summary of significant results for Hypothesis 5 |  |  |  |  |  |  |
| --- | --- | --- | --- | --- | --- | --- |
| Site | Question/Factor | Metric | LEBA item | Correlation coefficient<sup>1</sup> | p-value<sup>2</sup> | Interpretation |
| switzerland | I spend 30 minutes or less per day (in total) outside | PAT1000 | F2, 04 | -0.33 | 0.018 | The more often 30 minutes or less are spent outside, the shorter the period above 1000 lx |
| switzerland | I spend more than 3 hours per day (in total) outside | PAT1000 | F2, 07 | 0.46 | <0.001 | The more often >3 hours per day are spent outside, the longer the period above 1000 lx |
| switzerland | I spend more than 3 hours per day (in total) outside | TAT1000 | F2, 07 | 0.45 | <0.001 | The more often > 3 hours per day are spent outside, the higher the time above 1000 lx |
| switzerland | I spend as much time outside as possible | M10m | F2, 08 | 0.39 | 0.003 | The more spend time outside as possible, the brighter the brightest 10 hours (mean) |
| switzerland | I spend as much time outside as possible | PAT1000 | F2, 08 | 0.47 | <0.001 | The more spend time outside as possible, the longer the period above 1000 lx |
| switzerland | I spend as much time outside as possible | TAT1000 | F2, 08 | 0.48 | <0.001 | The more spend time outside as possible, the higher the time above 1000 lx |
| switzerland | I spend as much time outside as possible | TAT250 | F2, 08 | 0.32 | 0.011 | The more spend time outside as possible, the higher the time above 250 lx |
| switzerland | I look at my smartwatch within 1 hour before attempting to fall asleep | LLiT250 | F3, 13 | 0.34 | <0.001 | The more often smartwatches are looked at within 1h before attempting to fall asleep, the later the last time above 250 lx |
| switzerland | I use an alarm with a dawn simulation light | SCmelEDI | F5, 22 | -0.34 | <0.001 | The more an alarm with a dawn simulation light is used, the warmer the light colour temperature |
| switzerland | I turn on the lights immediately after waking up | FLiT250 | F5, 23 | -0.35 | 0.002 | The more often the lights are turned on immediately after waking up, the earlier the first time above 250 lx |
| <sup>1</sup> Spearman correlation coefficient between the metric and the LEBA question/factor |  |  |  |  |  |  |
| <sup>2</sup> p-values are adjusted for multiple comparisons using the false-discovery-rate for n= 168 comparisons |  |  |  |  |  |  |

## Discussion

In this observational study, we investigated light exposure and light exposure behaviour differences in 39 participants in their everyday lives over 30 consecutive days. Participants were recruited in Basel, Switzerland, and Kuala Lumpur, Malaysia, two locations that differ in culture, climate, urban environment and photoperiod. Given the multitude of available light metrics to quantify objective light exposure and a lack of standards which of those metrics are to be reported (Spitschan et al., 2019, 2023), we also investigated which specific light metrics differed between these sites (**RQ 1**).

### Objective light exposure is different between Switzerland and Malaysia beyond the differences in photoperiod

We found that participants in Switzerland spent significantly more time in bright levels likely corresponding to daylight (i.e., >1000 lux melanopic EDI) than participants in Malaysia during the day. Participants in Switzerland also stayed longer and about twice as often in recommended light levels during the day (i.e., >250 lux melanopic EDI). This is generally expected, given the longer photoperiod in Switzerland compared to the Malaysian site (see **Fig 1D**). However, these findings still hold when adjusting for the longer photoperiod in Switzerland, indicating differences between the sites. Furthermore, the brightest time of day was significantly brighter for participants in Switzerland compared to participants in Malaysia, with a difference of about 0.5 log_10_ units, indicating that participants in Switzerland experienced their brightest time of day with an intensity approximately 3.16 times higher. This potentially hints towards differences in behaviour since theoretical access to daylight based on photoperiod was comparable, although shorter photoperiods also offer lower daylight levels. A more likely explanation might be, though, that in both sites, melanopic EDI levels during the day indicate predominantly indoor environments. Still, the significantly higher and longer exposure to bright light in participants in Switzerland suggests better access to daylight while being indoors, compared to Malaysia.

As expected, the last time of day with bright light exposure above 250 lux melanopic EDI was later by about 1.5 hours in Switzerland compared to Malaysia, which can partially be explained by the later dusk in Switzerland (about 0.8 hours), but also by daylight savings time (DST), which was in place in Switzerland during data collection, but not Malaysia. This means that participants in Switzerland experienced light until later in their day, which could contribute to later chronotypes (although the reverse could also be true). Previous studies have shown an interaction of chronotype and light exposure as well as sleep timings and light exposure (Refinetti, 2019; Wahl et al., 2019; Wright et al., 2013; Zerbini et al., 2020, 2021). Unfortunately, chronotype was not formally assessed in this study, but based on PSQI-derived mid-sleep (MS) as a proxy for chronotype, we also found a correlation between chronotype and mean timing of light above 250 mel EDI in Malaysia (r=0.22, *p<0.001*) but not in Switzerland (r=0.05, *p=0.27*). Note that we did not categorise chronotype into “early”, “late”, or “intermediate” types (Roenneberg et al., 2019) since this type of categorisation is not particularly meaningful in our case, spanning two very distinct geographic locations and cultures. According to this binning, mid-sleep before 3 a.m. is “early”, while mid-sleep after 4 a.m. is “late” (while both early and late have sub-categories such as “moderate” or “definite”). Applying these categories across sites would reduce the difference to a significantly higher proportion of early types in Switzerland compared to Malaysia. This ignores several important distinctions that are retained in the continuous scale: the distribution of chronotypes between sites (Fig. S5, and Fig. 2) shows a clear difference in the median but also scale. Except for one particularly late participant in Switzerland, Swiss participants do not exhibit a large variance compared to Malaysia. While the phase of the photoperiod is different between the sites (e.g., based on solar noon), explaining about one fifth of the difference in the mean, other factors must contribute to this difference and the wider distribution as well. For example, a strong daytime stimulus of natural daylight and low levels of artificial light at night have been shown to both reduce the variance of melatonin rhythms (which are also used as a proxy for chronotype), and result in a shift towards earlier rhythms, i.e. earlier chronotypes (Stothard et al., 2017; Wright et al., 2013).

Participants in Switzerland in our study did also avoid light exposure above 10 lux mel EDI significantly earlier than participants in Malaysia by about 1 hour and 10 minutes (i.e., the timing of brighter light exposure in the evening). This means that although they still experienced some light later in their 24-day, it was dimmer than the light participants in Malaysia experienced before their bedtime. Additionally, participants in Switzerland averaged about 5 times (∼0.6 log_10_ units) lower melanopic EDI levels during evenings, suggesting that participants in Switzerland seem to have actively avoided bright light exposure or preferred dimly lit environments. The latter could be explained by cultural differences in light preference, with a potential preference for brighter and whiter electric light in Asian cultures (Baniya et al., 2015). We cannot provide evidence beyond speculation here as we did not assess participants’ lighting preferences.

### Objective light exposure patterns reveal strong differences between individuals within the sites but do not follow recommendations for healthy light exposure in either site

Our analysis of light exposure patterns reveals that while, on average, the above-mentioned differences in light exposure hold up, the explanatory power of a single exposure pattern per site is negligible. Instead, each site consists of several individual patterns, with a within-site variation greater than between sites. This is encouraging, as it means people on an individual level have influence over their light exposure pattern beyond their environmental or cultural baseline. Thus, interventions to modify personal light exposure patterns might have a greater chance of success than they would have with very similar individual patterns across each site. While we did not assess socioeconomic status (SES), it should be noted that Malaysia and Switzerland show different ranges in GDP per capita, differing by almost a factor of 10 in 2024 (Malaysia: $11,867.3 vs. Switzerland: $103,669.9) (World Bank Group, n.d.). This is important because, although there is no systematic data on how SES might affect light exposure, there are indications that light exposure at night depends on SES (Crea, 2021; Nadybal et al., 2020).

While light exposure was more favourable in the Swiss dataset, neither participants in Switzerland nor Malaysia spent much of their time in bright light above 250 lux melanopic EDI, as recommended by Brown and colleagues (Brown et al., 2022). For participants in Switzerland, it is about 12 minutes per hour of daylight, and 7 minutes for participants in Malaysia, i.e., 20% and 12% of the available daytime, respectively, which corroborates previous findings where participants in Switzerland spent only 14% of their light logger wearing time above 250 lux melanopic EDI during daytime (Stefani et al., 2024). In the evening, the participants in Switzerland in our study reduced their light exposure level earlier to the recommended 10 lux melanopic EDI or below. Still, both sites had times of elevated light exposure levels after dusk.

### Self-reported light exposure behaviour does not differ between Switzerland and Malaysia and is stable over time

A second goal of this study was to understand if subjectively reported light exposure behaviour measured with the LEBA instrument showed cultural differences between the two locations (**RQ 2**). Since LEBA is a new instrument, cultural differences, also in specific items or scores of the questionnaire, are yet to be investigated and tested. In contrast to our expectations, LEBA items and factors did not show a significant difference between Malaysia and Switzerland, meaning that light exposure behaviour assessed with the LEBA was comparable across these cultures, at least in our sample. Due to the timeframe overlap in the Malaysian dataset (i.e., participants were asked every few days about the previous *four* weeks), we were also able to investigate if items and scores were stable over time, which we confirmed for most of the LEBA items. All 23 questions and 1 out of 5 factors did not vary significantly in more than 50% of participants. While the other factors showed variation over time, it was very small, with an average sum of 3-to-4-point total differences across nine survey periods. Unfortunately, we were unable to do this analysis in the Swiss dataset because the timespans used for the instruction in the LEBA questionnaires were not comparable. Therefore, we cannot be sure if the stability of LEBA items can be generalised beyond the Malaysian site, but we would hypothesise so based on the overall lack of differences between cultures in LEBA items and scores.

### Self-reported light exposure behaviour is reflected in objective light exposure but only in participants in Switzerland

Finally, we wanted to test if subjectively reported light exposure behaviour can be related to objectively measured light exposure and, again, if these relationships might depend on location (Research Question 3). The results clearly show that there are strong correlations between certain light exposure metrics and answers to the LEBA questionnaire, but only in the Swiss dataset. After adjustment for multiple comparisons, none of the correlations in the Malaysian dataset remained significant. Interestingly, the average (absolute) correlation was stronger in the Malaysia site than in Switzerland, indicating a higher variance in either light exposure metrics or LEBA answers in the Malaysia dataset compared to Switzerland. For the Swiss dataset, ten of the hypothesised correlations were shown to be significant after adjustment with an average medium effect size (Cohen, 1988, 1992).

We acknowledge that differences in the sites might also be due to the different phrasing of the LEBA: whereas it questioned participants about the *past few days* in Switzerland, it asked about the *past four weeks* in Malaysia. Thus, the differences in correlative strength might also be due to participants being able to recollect light exposure-related behaviour better for a few days compared to the last month.

### Strength and weaknesses

Our study has several strengths and weaknesses. We sampled data from a relatively small cohort of n=19 and n=20 per site, respectively, without controlling for SES, body mass index (BMI) or mental/psychiatric disorders, which were previously shown to alter light exposure (Badpa et al., 2023; Crea, 2021; Deprato et al., 2025; Gubin et al., 2023, 2025; Nadybal et al., 2020) However, given the longitudinal data of four weeks per participant, we are confident that we have enough data points to rely on the model results that we presented. Since we only collected data from one site per country, we cannot be sure that these sites represent the entire country, but this was also not the overarching research goal. Unfortunately, we lacked information on chronotype, weekday type information (work day vs. free day) as well as type of activity or work schedules, so we could not compute statistics that would have included such information despite previous studies showing their potential influence (Biswas et al., 2025). The light logger we used, the SPECCY wearable, also did not record activity (actimetry data) to save battery, so we could not use this information to detect the non-wear time of the device nor estimate chronotype based on activity. We have, however, approximated chronotype based on mid-sleep from the PSQI questionnaire.

Our study, though, adds to the novel literature on light exposure *behaviour* and is the first to relate subjectively assessed light behaviour to objectively recorded light exposure behaviour. Our data sampling is both longitudinal and takes place at high frequency, allowing for fine-grained and highly reliable data analysis. By positioning the light logger at chest level, we were able to better approximate physiologically relevant light exposure than at wrist positions. Finally, we add to the growing literature on individual and cultural differences, which need to be factored in for better and fine-tuned recommendations and provide here a dataset that generated novel hypotheses that can be tested explicitly in the future.

## Conclusion

This study offers a detailed investigation into the differences in light exposure and light exposure behaviour between two distinct geographical and cultural contexts: Switzerland and Malaysia. Key findings reveal significant differences in objective light exposure between the two sites. Participants in Switzerland experienced brighter and longer daylight exposure, even when accounting for photoperiod differences while avoiding evening light exposure earlier and to a greater extent than participants in Switzerland. These results suggest behavioural and cultural preferences or differences in access to daylight, all of which influence individual light exposure patterns. Additionally, our finding that subjective light exposure behaviour correlates with objective light metrics only in the Swiss dataset underscores potential cultural or methodological differences in self-reporting accuracy. Despite this, the LEBA instrument proved stable over time, offering preliminary evidence for its reliability.

While our study advances the understanding of light exposure behaviours and cultural nuances, it also highlights several areas for future exploration. Investigating preferences for electrical light colour and intensity could also clarify the cultural differences hypothesised in this study. Expanding the sample size and geographic scope would improve the generalisability of these findings, while further validation of the LEBA instrument in diverse populations is essential to confirm its cross-cultural applicability and whether it could be used as an approximation for objectively assessed light exposure. Similarly, we recommend that future studies include SES as confounding variable and even systematically study differences in light exposure based on SES (Spitschan & Joyce, 2023).

From an applied perspective, these findings have implications for designing culturally informed light exposure guidelines to promote healthier circadian alignment. The apparent avoidance of evening light exposure in the participants in Switzerland aligns with recommendations for better sleep and circadian health. In contrast, the ‘preferences’, or at least exposure patterns, observed among the participants in Malaysia highlight the need for interventions aimed at reducing evening light exposure. These could include promoting the use of warmer, dimmer lighting in the evening and adjusting the timing of brighter light exposure during the day. Such strategies should consider local preferences and location-specific constraints, such as Malaysia’s daytime heat, which often discourages outdoor activities and exposure during daylight hours. Future studies should explore how environmental and behavioural interventions could optimise light exposure for health outcomes across diverse populations and climates to be feasible in the real world beyond Western locations and in light of global warming.

In conclusion, this study provides a foundation for understanding the interaction of light exposure behaviour, culture, and geography, opening new avenues for research and intervention strategies. By addressing the identified limitations and building on the strengths of our approach, future research can continue to uncover actionable insights into the critical role of light exposure in human circadian and sleep health.

## Data availability statement

Data and code are available on https://github.com/tscnlab/BillerEtAl_JExpoSciEnvironEpidemiol_2025 and are archived on Zenodo (10.5281/zenodo.16872096) under a permissive CC-BY 4.0 Attribution license.

The webpage https://tscnlab.github.io/BillerEtAl_JExpoSciEnvironEpidemiol_2025/ contains a Notebook of code and code outputs.

## Contributions

Conceptualisation: AMB, CC, MS, VK, JZ

Data curation: JZ

Formal analysis: JZ

Funding acquisition: CC, MS, OS, VK

Methodology: AM, LR, MG, MS, MYS, OS, VK

Project administration: CC, OS, VK

Software: JZ Resources: CC, VK

Supervision: CC, MS, OS, VK

Validation: -

Visualisation: JZ

Writing – original draft: AMB, JZ

Writing – review & editing: AMB, AM, CC, LR, MG, MS, MYS, OS, VK, JZ

## Acknowledgements

We thank ETH Zurich and Leading House Asia for awarding the RPG Research Partnership Grant ASEAN Number 072022 (“Shaping global human light behaviour – from chaos to order”) to OS and VK.

## Ethical approval

At the Swiss site, the study was conducted according to the Declaration of Helsinki and the principles of Good Clinical Practice and approved by the Ethikkommission Nordwest-und Zentralschweiz (EKNZ) under Basec ID # 2023-00827 on 4^th^ July 2023.

Data collection at the Malaysian site was approved by Monash University Human Research Committee (MUHREC, Project ID: 39378).

## Supplementary material

### Supplementary Tables

**Table S1.** Scoring of Light Exposure Behaviour Assessment (LEBA) items. Note that R denotes items that are reversed-scored. F1-F5 denotes the calculated Factors.

| Factor name | Score |
| --- | --- |
| F1: Wearing blue light filters | 01+02+03 |
| F2: Spending time outdoors | 04(R)+05+06+07+08+09 |
| F3: Using phone and smartwatch in bed | 10+11+12+13+14 |
| F4: Using light before bedtime | 15+16+17+18 |
| F5: Using light in the morning and during daytime | 19+20+21+22+23 |

**Table S2.** Scoring of the Pittsburgh Sleep Quality Index (PSQI) items.

| **Component** | **Question (Q)** | **Component Score** |
| --- | --- | --- |
| **(1)** Subjective sleep quality | #6 | 0-3 |
| **(2)** Sleep latency | #2<br>#5a | Sum of Q2 (0-3)+Q5a (0-3):<br>If 0 = 0<br>1-2 = 1<br>3-4 = 2<br>5-6 = 3 |
| **(3)** Sleep duration | #4 | 0-3 |
| **(4)** Habitual sleep efficiency<br>(Sleep efficiency = hours slept / hours in bed) X 100%) | #1<br>#3<br>#4 | If >85% = 0<br>75-84% = 1<br>65-74% = 2<br><65% = 3 |
| **(5)** Sleep disturbance | #5b-5j | Sum of Q5b (0-3) to Q5 (0-3):<br>If 0 = 0<br>1-9 = 1<br>10-18 = 2<br>19-27 = 3 |
| **(6)** Use of sleep medication | #7 | 0-3 |
| **(7)** Daytime dysfunction | #8<br>#9 | Sum of Q8 (0-3) +Q9 (0-3):<br>If 0 = 0<br>1-2 = 1<br>3-4 = 2<br>5-6 = 3 |
| **Global PSQI Score** |  | **Sum of Components 1-7** |

**Table S3.**
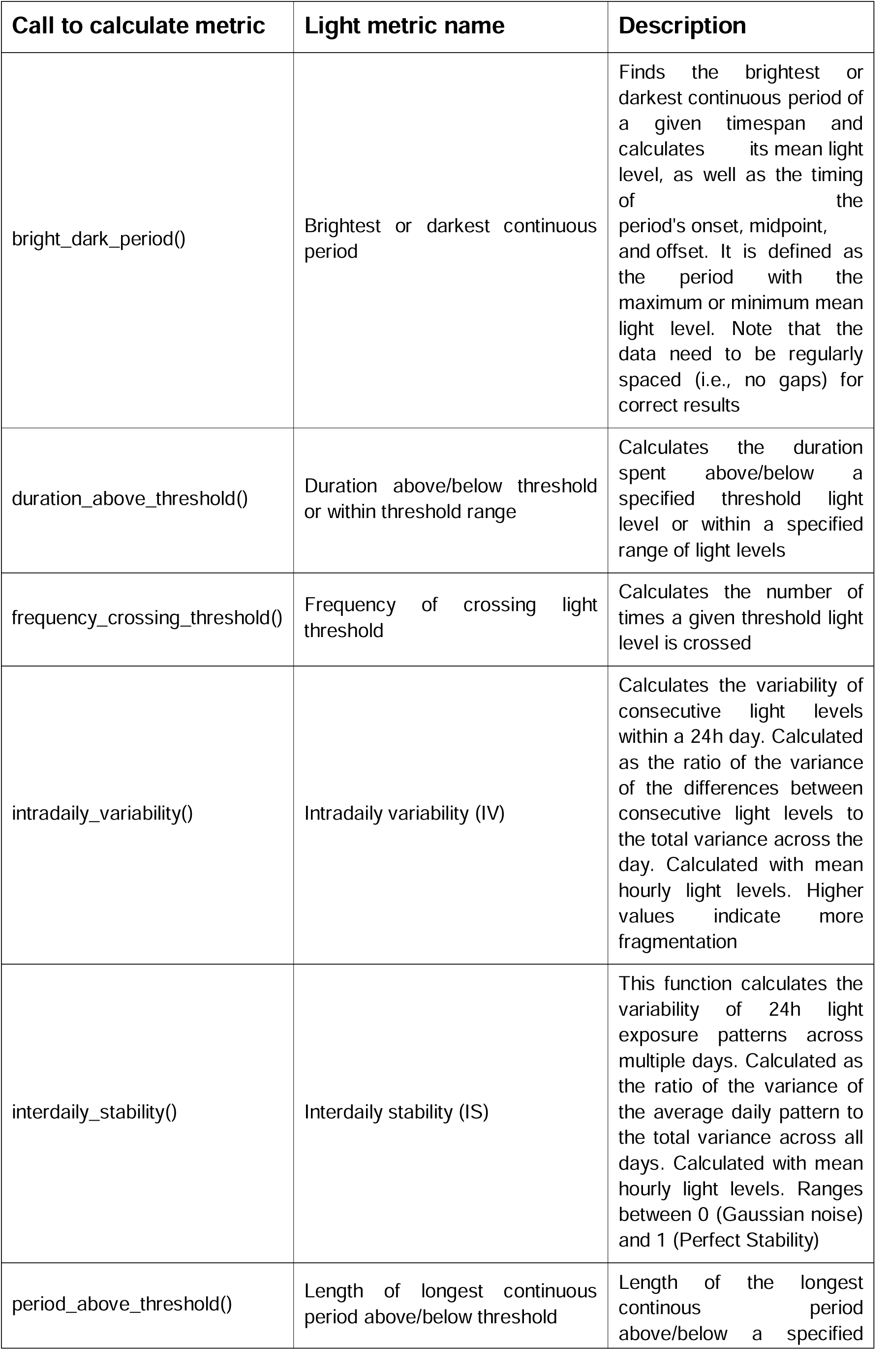
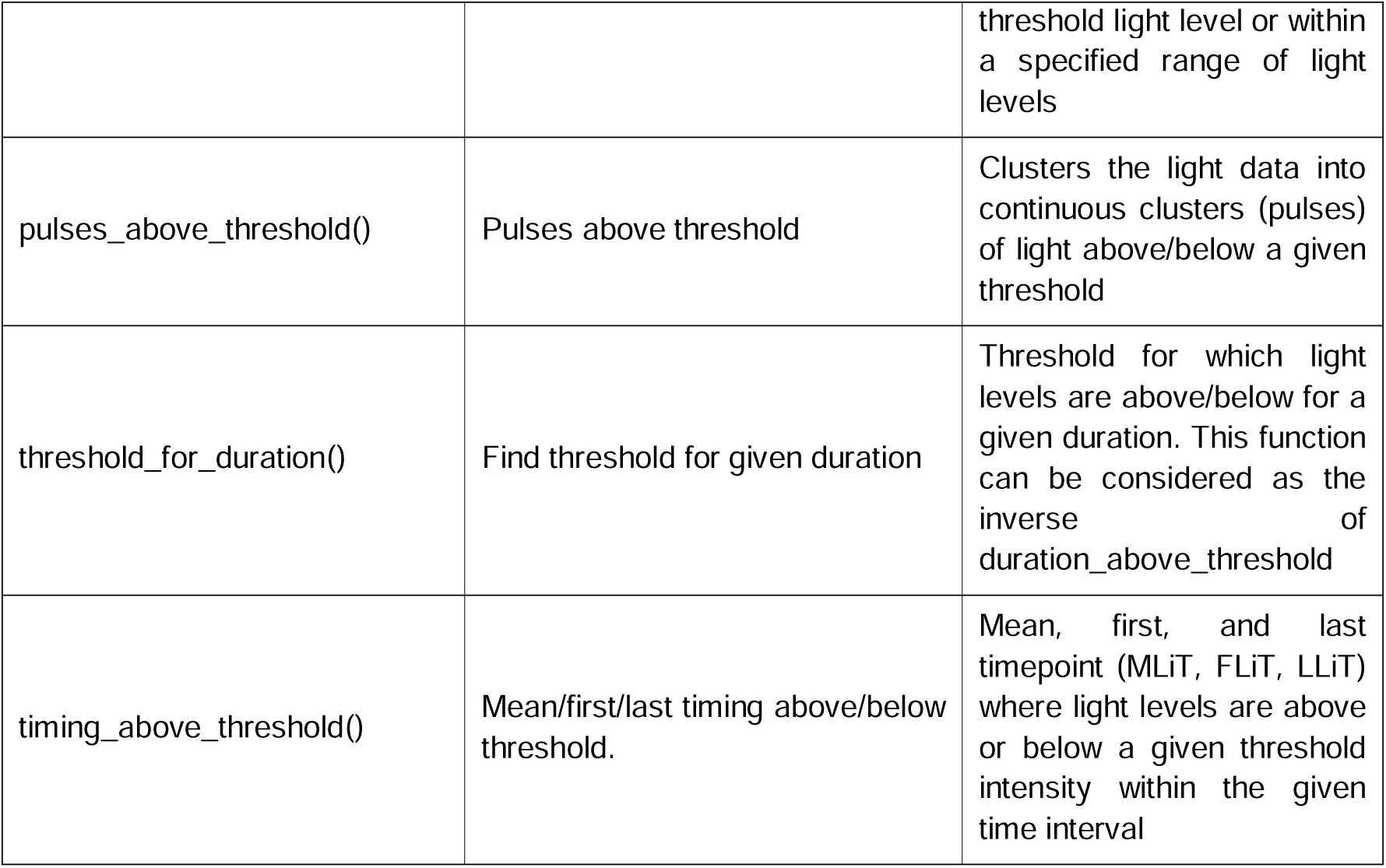
Overview of objectively derived outcome variables from light logging data and their calculations.

**Table S4.**
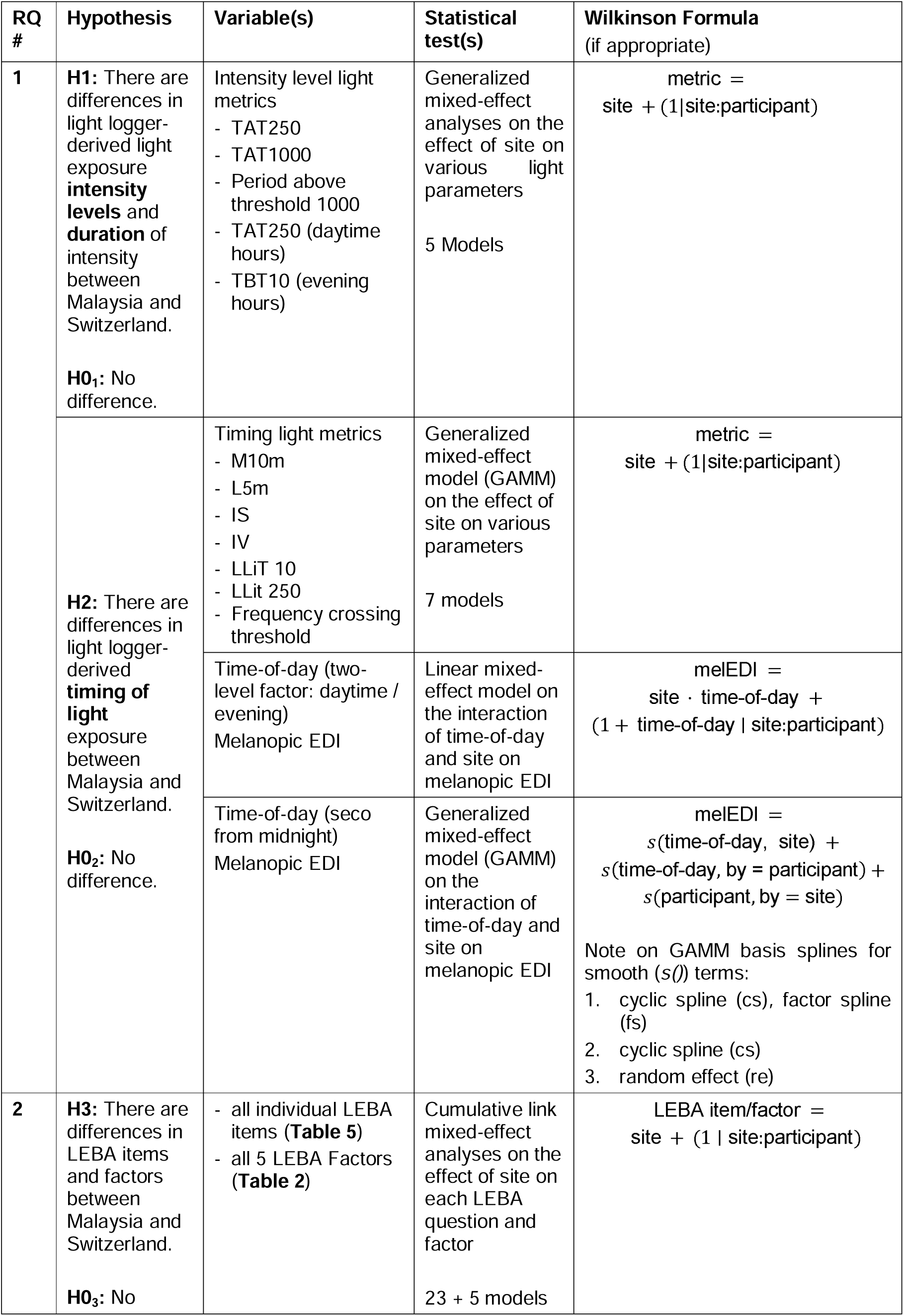
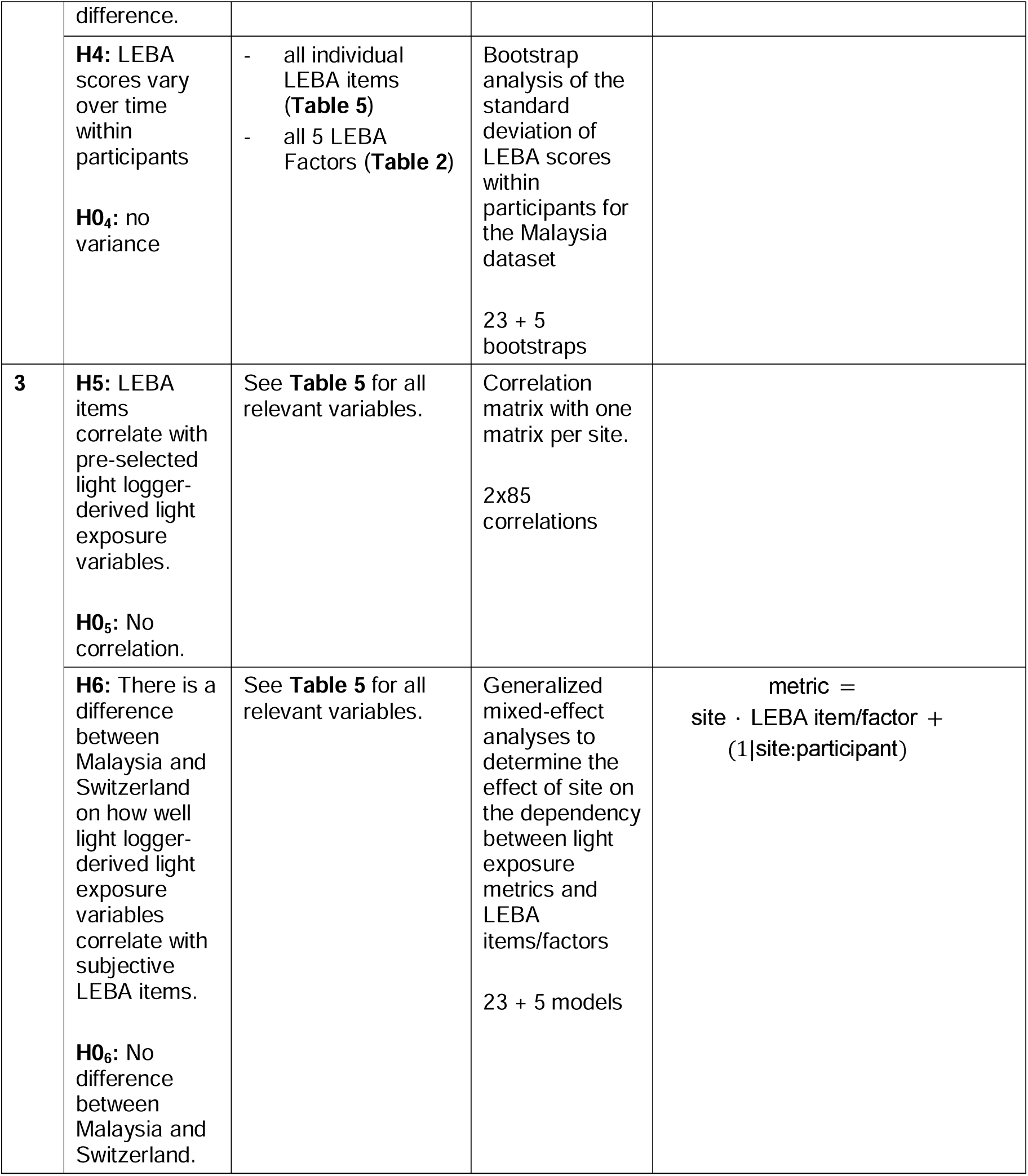
Overview of research questions, hypotheses and statistical tests to address the hypotheses. Abbreviations: GAMM, Generalised additive mixed-effect model; H0_x_, Null hypothesis; HX, Hypothesis X; IS, Interdaily Stability; IV, Intradaily variability; LEBA, Light Exposure Behaviour Assessment; LLiT, Last time above threshold; LLiT 10, Low light intensity threshold at 10 lux; LLiT 250, Low light intensity threshold at 250 lux; L5m, Mean across darkest 5 hours; M10m, Mean across brightest 10 hours; melEDI, Melanopic equivalent daylight illuminance; RQ, Research question; TAT250, Time above threshold 250 lux; TAT1000, Time above threshold 1000 lux; TBT10, Time below threshold 10 lux.

**Table S5.**
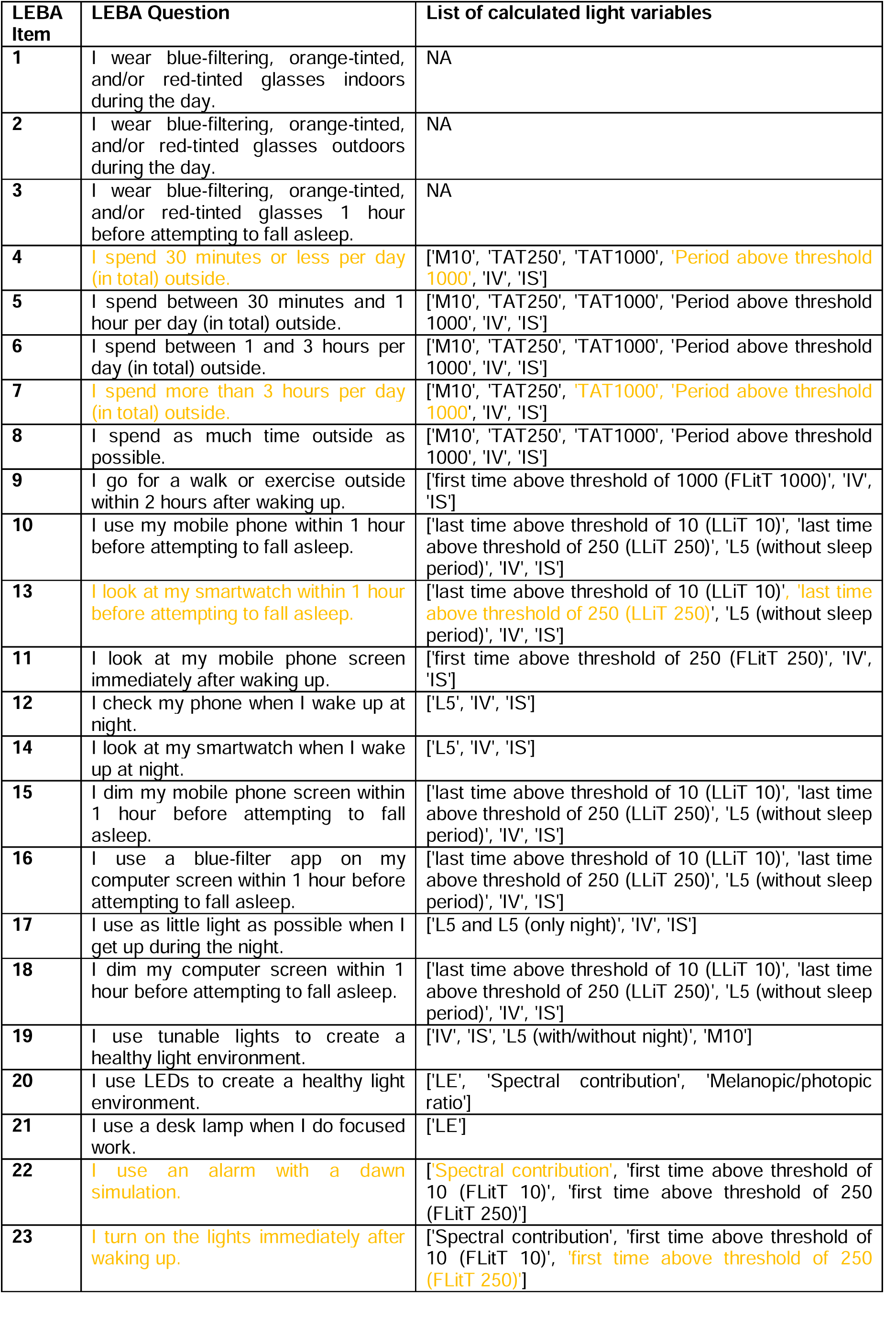
List of Light Exposure Behaviour Assessment (LEBA) items and corresponding objectively derived light variables from light logging data. In yellow are the hypothesised and significant relationships in the Swiss dataset. Abbreviations: FLiT 10, First time above threshold of 10 lux; FLiT 250, First time above threshold of 250 lux; FLiT 1000, First time above threshold of 1000 lux; IS, Interdaily Stability; IV, Intradaily Variability; LE, Light Exposure; L5, Mean across darkest 5 hours; L5 (only night), Mean across darkest 5 hours during the night; LLiT 10, Last time above threshold of 10 lux; LLiT 250, Last time above threshold of 250 lux; M10, Mean across brightest 10 hours; TAT250: Time above threshold of 250 lux; TAT1000: Time above threshold of 1000 lux.

**Table S6.** Overview of missing data for the two datasets. A, Shows the overview for Malaysia and B, for Switzerland. Abbreviations: Min, minimum; Max, maximum.

|  | min | max | median | mean | total |
| --- | --- | --- | --- | --- | --- |
| Number of gaps | 0 | 7 | 0 | 1 | 22 |
| Duration missed by available data points | 0 days | 14 days | 0 days | 1 day | 29 days |
| Duration covered by available data points | 11 days | 30 days | 30 days | 26 days | 509 days |
| Number of missing data points | 0 | 21,040 | 0 | 2,200 | 41,797 |
| Number of available data points | 16,604 | 43,996 | 43,211 | 38,627 | 733,907 |
| Percentage of missing data points | 0.0% | 48.7% | 0.0% | 5.4% | 5.4% |
| min, max, median, mean, and total values for 19 participants |  |  |  |  |  |

|  | min | max | median | mean | total |
| --- | --- | --- | --- | --- | --- |
| Number of gaps | 1 | 4 | 1 | 2 | 35 |
| Duration missed by available data points | <1 day | 11 days | <1 day | 1 day | 28 days |
| Duration covered by available data points | 7 days | 30 days | 29 days | 27 days | 554 days |
| Number of missing data points | 21 | 16,017 | 44 | 2,035 | 40,698 |
| Number of available data points | 11,476 | 44,586 | 42,974 | 39,954 | 799,070 |
| Percentage of missing data points | 0.0% | 40.4% | 0.1% | 5.9% | 4.8% |
| min, max, median, mean, and total values for 20 participants |  |  |  |  |  |

**Table S7.**
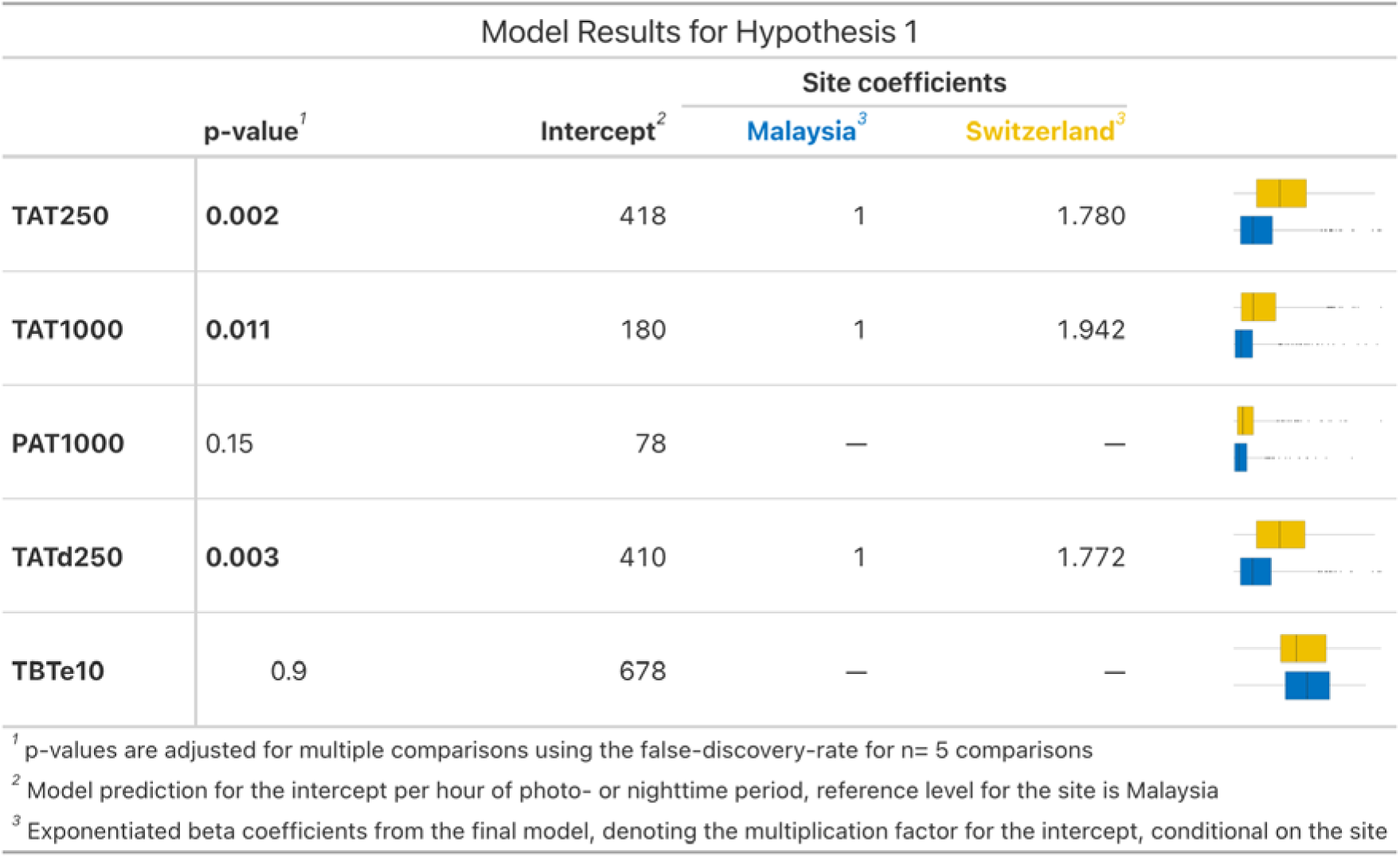
Model results for H1. P-values are considered significant below 0.05. Abbreviations: PAT1000, Period above threshold 1000, TAT250, Time above threshold of 250 lux; TAT1000, Time above threshold of 1000 lux; TATd250, Time above dynamic threshold 250 lux; TBTe10, Time below threshold 10 lux.

| Model Results for Hypothesis 1 |  |  |  |  |
| --- | --- | --- | --- | --- |
|  | p-value<sup>1</sup> | Intercept<sup>2</sup> | Site coefficients |  |
|  |  |  | Malaysia<sup>3</sup> | Switzerland<sup>3</sup> |
| TAT250 | 0.002 | 418 | 1 | 1.780 |
| TAT1000 | 0.011 | 180 | 1 | 1.942 |
| PAT1000 | 0.15 | 78 | — | — |
| TATd250 | 0.003 | 410 | 1 | 1.772 |
| TBTd10 | 0.9 | 678 | — | — |

<sup>1</sup> p-values are adjusted for multiple comparisons using the false-discovery-rate for n= 5 comparisons
<sup>2</sup> Model prediction for the intercept per hour of photo- or nighttime period, reference level for the site is Malaysia
<sup>3</sup> Exponentiated beta coefficients from the final model, denoting the multiplication factor for the intercept, conditional on the site

**Table S8.**
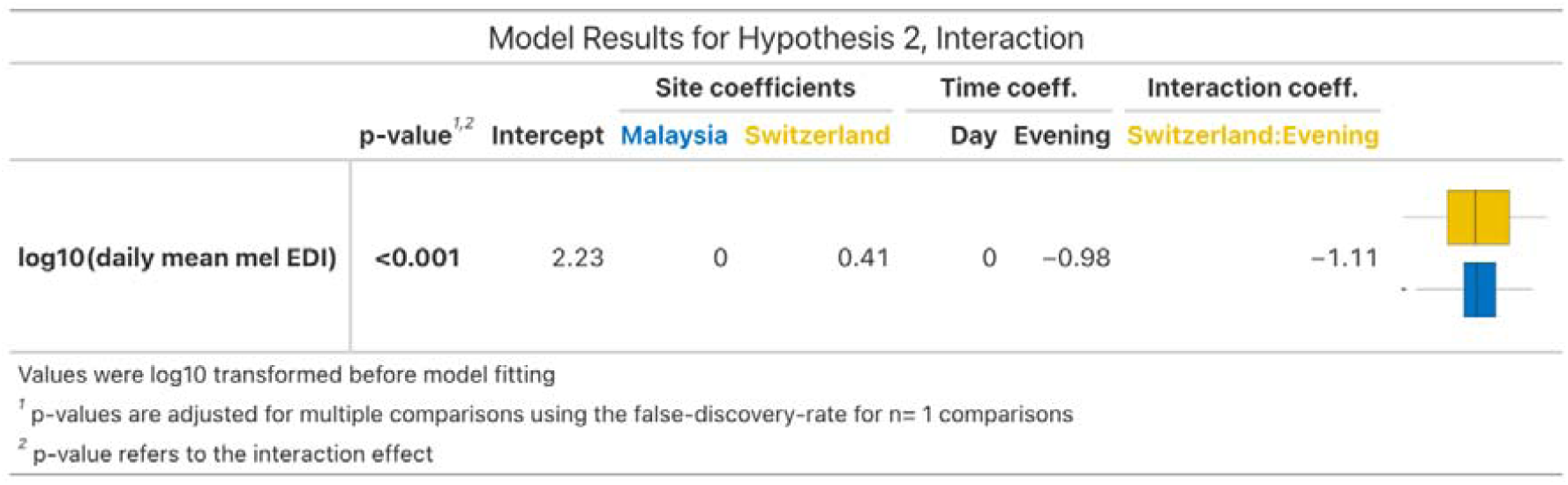
Model results for H2. P-values are considered significant below 0.05. Abbreviations: Mel EDI, melanopic equivalent daylight illumination.

| Model Results for Hypothesis 2, Interaction |  |  |  |  |  |  |  |
| --- | --- | --- | --- | --- | --- | --- | --- |
|  | p-value<sup>1,2</sup> | Intercept | Site coefficients |  | Time coeff. |  | Interaction coeff. |
|  |  |  | Malaysia | Switzerland | Day | Evening | Switzerland:Evening |
| log10(daily mean mel EDI) | <0.001 | 2.23 | 0 | 0.41 | 0 | -0.98 | -1.11 |
| Values were log10 transformed before model fitting |  |  |  |  |  |  |  |
| <sup>1</sup> p-values are adjusted for multiple comparisons using the false-discovery-rate for n= 1 comparisons |  |  |  |  |  |  |  |
| <sup>2</sup> p-value refers to the interaction effect |  |  |  |  |  |  |  |

**Table S9.**
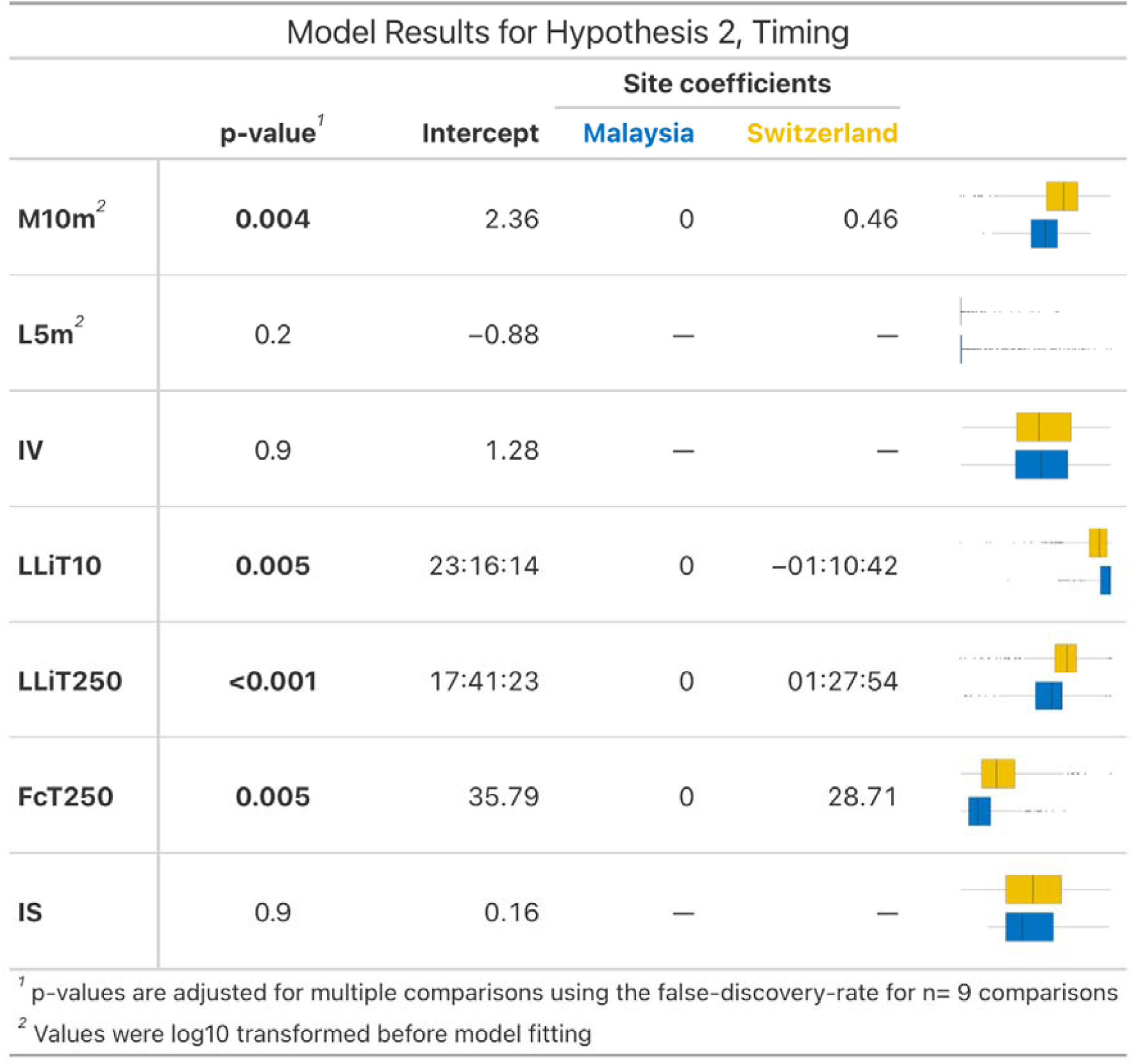
Model results for H2. P-values are considered significant below 0.05. Abbreviations: FcT250, Frequency of crossing light threshold at 250 lux; FLiT 10, First time above threshold of 10 lux; FLiT 250, First time above threshold of 250 lux; FLiT 1000, First time above threshold of 1000 lux; IS, Interdaily Stability; IV, Intradaily Variability; L5m, Mean across darkest 5 hours; LE, Light Exposure; LLiT 10, Last time above threshold of 10 lux; LLiT 250, Last time above threshold of 250 lux; M10m, Mean across brightest 10 hours;

| Model Results for Hypothesis 2, Timing |  |  |  |  |
| --- | --- | --- | --- | --- |
|  | p-value<sup>1</sup> | Intercept | Site coefficients |  |
|  |  |  | Malaysia | Switzerland |
| M10m<sup>2</sup> | 0.004 | 2.36 | 0 | 0.46 |
| L5m<sup>2</sup> | 0.2 | -0.88 | — | — |
| IV | 0.9 | 1.28 | — | — |
| LLiT10 | 0.005 | 23:16:14 | 0 | -01:10:42 |
| LLiT250 | <0.001 | 17:41:23 | 0 | 01:27:54 |
| FcT250 | 0.005 | 35.79 | 0 | 28.71 |
| IS | 0.9 | 0.16 | — | — |

<sup>1</sup> p-values are adjusted for multiple comparisons using the false-discovery-rate for n= 9 comparisons
<sup>2</sup> Values were log10 transformed before model fitting

**Table S10.**
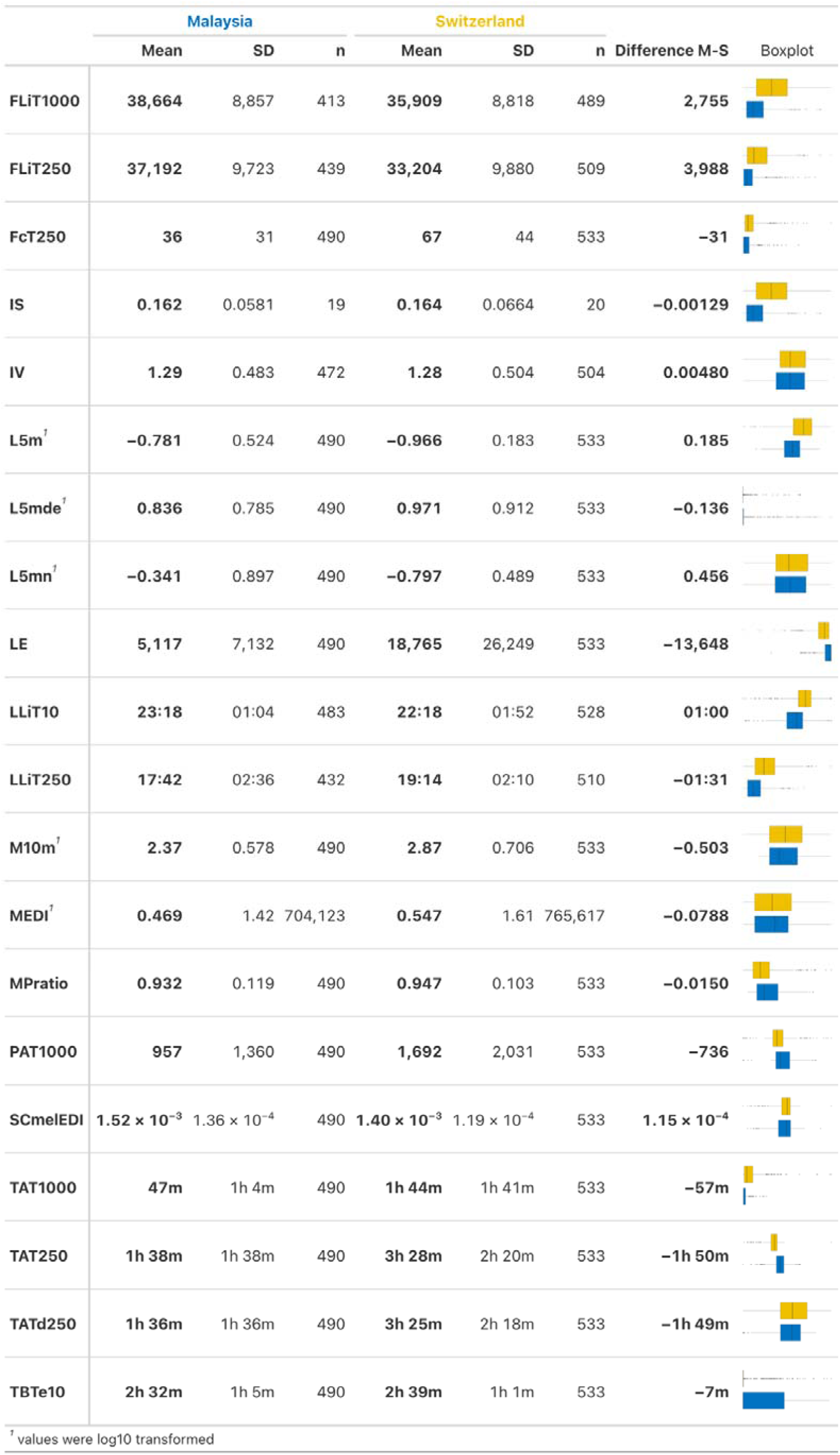
Differences in descriptive statistics for light exposure metrics between Malaysia and Switzerland. Mean, standard deviation (SD), sample size (n) and mean differences between the two locations (M-S) are presented for each variable. Abbreviations: FcT250, Frequency of crossing light threshold at 250 lux; FLiT250, First time above threshold of 250 lux; FLiT1000, First time above threshold of 1000 lux; IS, Interdaily Stability; IV, Intradaily Variability; LE, Light Exposure; L5m, Mean across darkest 5 hours; L5mde, Mean duration of exposure during the least active 5 hours; L5mn, Number of exposures during the least active 5 hours; LLiT10, Last time above threshold of 10 lux; LLiT250, Last time above threshold of 250 lux; M10m, Mean across brightest 10 hours; mel EDI, melanopic equivalent daylight illuminance; MPratio, Ratio between melanopic and photopic illuminance; PAT1000, Period above threshold of 1000 lux; SCmelEDI, Spectal composition equivalent daylight illuminance; TAT250, Time above threshold of 250 lux; TAT1000, Time above threshold of 1000 lux; TATd250, Time above dynamic threshold of 250 lux; TBTe10, Time below threshold of 10 lux.

**Table S11.** Stability of LEBA items and factors over time. For a description of each LEBA item, see Table S5.

| Hypothesis 4: Extent of stable LEBA items and factors over time |
| --- |
| **stable over time in 75-100% of participants** |
| Factor 1: Item 01, Factor 1: Item 02, Factor 1: Item 03, Factor 2: Item 09, Factor 3: Item 10, Factor 3: Item 13, Factor 3: Item 14, Factor 5: Item 19, Factor 5: Item 21, Factor 5: Item 22, Factor 1 |
| **stable over time in 50-75% of participants** |
| Factor 2: Item 04, Factor 2: Item 05, Factor 2: Item 06, Factor 2: Item 07, Factor 2: Item 08, Factor 3: Item 11, Factor 3: Item 12, Factor 4: Item 15, Factor 4: Item 16, Factor 4: Item 17, Factor 4: Item 18, Factor 5: Item 20, Factor 5: Item 23 |
| **stable over time in 25-50% of participants** |
| Factor 3 |
| **stable over time in 25-0% of participants** |
| Factor 2, Factor 4, Factor 5 |

**Table S12.**
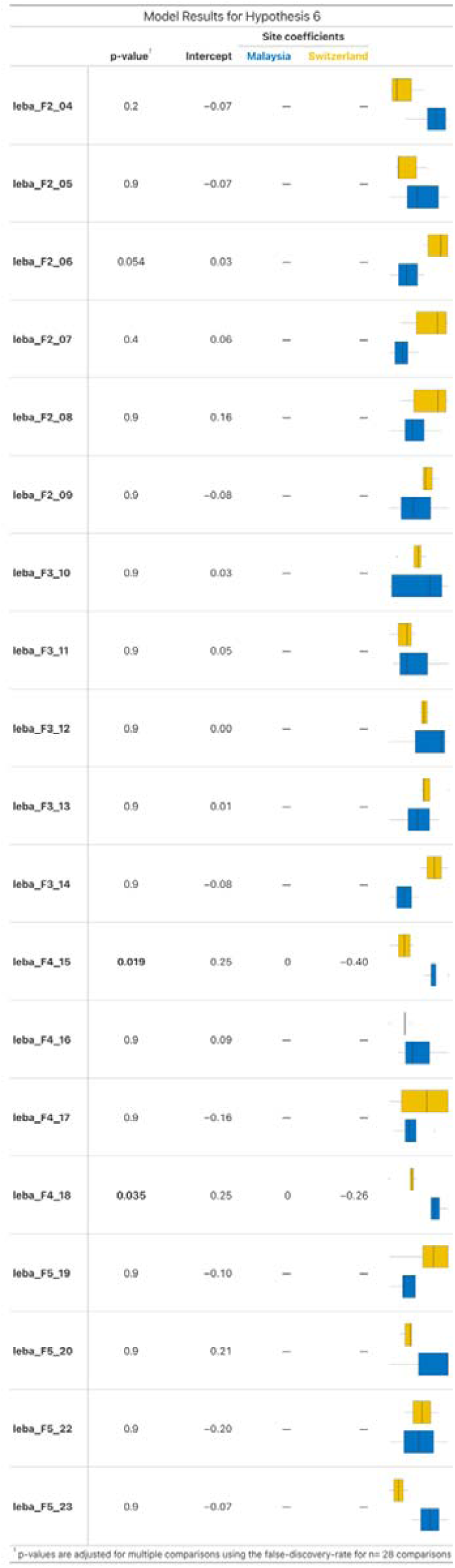
Model results of H6. P-values are considered significant below 0.05. For a description of each LEBA item, see Table S5.

**Table S13.** Pittsburgh Sleep Quality Index comparisons between Malaysia and Switzerland at Day 0. P-values are considered significant below 0.05.

| Characteristic | Malaysia<br>N = 19<sup>1</sup> | Switzerland<br>N = 20<sup>1</sup> | p-value<sup>2</sup> |
| --- | --- | --- | --- |
| Subjective sleep quality |  |  | >0.9 |
| 0 | 3 (16%) | 3 (15%) |  |
| 1 | 15 (79%) | 16 (80%) |  |
| 2 | 1 (5.3%) | 1 (5.0%) |  |
| 3 | 0 (0%) | 0 (0%) |  |
| Sleep latency |  |  | >0.9 |
| 0 | 7 (37%) | 6 (30%) |  |
| 1 | 9 (47%) | 10 (50%) |  |
| 2 | 3 (16%) | 3 (15%) |  |
| 3 | 0 (0%) | 1 (5.0%) |  |
| Sleep duration |  |  | 0.2 |
| 0 | 4 (21%) | 9 (45%) |  |
| 1 | 13 (68%) | 11 (55%) |  |
| 2 | 1 (5.3%) | 0 (0%) |  |
| 3 | 1 (5.3%) | 0 (0%) |  |
| Habitual sleep efficiency |  |  | 0.020* |
| 0 | 18 (95%) | 12 (60%) |  |
| 1 | 1 (5.3%) | 8 (40%) |  |
| 2 | 0 (0%) | 0 (0%) |  |
| 3 | 0 (0%) | 0 (0%) |  |
| Sleep disturbance |  |  | 0.5 |
| 0 | 2 (11%) | 2 (10%) |  |
| 1 | 17 (89%) | 16 (80%) |  |
| 2 | 0 (0%) | 2 (10%) |  |
| 3 | 0 (0%) | 0 (0%) |  |
| Use of sleep medication |  |  | >0.9 |
| 0 | 18 (95%) | 17 (85%) |  |
| 1 | 1 (5.3%) | 2 (10%) |  |
| 2 | 0 (0%) | 1 (5.0%) |  |
| 3 | 0 (0%) | 0 (0%) |  |
| Daytime dysfunction |  |  | 0.4 |
| 0 | 3 (16%) | 7 (35%) |  |
| 1 | 11 (58%) | 9 (45%) |  |
| 2 | 4 (21%) | 4 (20%) |  |
| 3 | 1 (5.3%) | 0 (0%) |  |
| Global PSQI | 5.00 (2.00,9.00) | 4.00 (1.00,10.00) | 0.8 |

<sup>1</sup> n (%); Median (Min,Max)
<sup>2</sup> \*p<0.05; \*\*p<0.01; \*\*\*p<0.001

**Table S14.** Pittsburgh Sleep Quality Index comparisons between Malaysia and Switzerland on Day 31. P-values are considered significant below 0.05.

| Characteristic | Malaysia<br>N = 19<sup>1</sup> | Switzerland<br>N = 20<sup>1</sup> | p-value<sup>2</sup> |
| --- | --- | --- | --- |
| Subjective sleep quality |  |  | 0.8 |
| 0 | 2 (11%) | 3 (17%) |  |
| 1 | 13 (68%) | 13 (72%) |  |
| 2 | 4 (21%) | 2 (11%) |  |
| 3 | 0 (0%) | 0 (0%) |  |
| missing | 0 | 2 |  |
| Sleep latency |  |  | 0.7 |
| 0 | 8 (42%) | 5 (28%) |  |
| 1 | 7 (37%) | 9 (50%) |  |
| 2 | 2 (11%) | 3 (17%) |  |
| 3 | 2 (11%) | 1 (5.6%) |  |
| missing | 0 | 2 |  |
| Sleep duration |  |  | 0.2 |
| 0 | 8 (42%) | 13 (72%) |  |
| 1 | 8 (42%) | 5 (28%) |  |
| 2 | 2 (11%) | 0 (0%) |  |
| 3 | 1 (5.3%) | 0 (0%) |  |
| missing | 0 | 2 |  |
| Habitual sleep efficiency |  |  | >0.9 |
| 0 | 16 (84%) | 17 (94%) |  |
| 1 | 2 (11%) | 1 (5.6%) |  |
| 2 | 1 (5.3%) | 0 (0%) |  |
| 3 | 0 (0%) | 0 (0%) |  |
| missing | 0 | 2 |  |
| Sleep disturbance |  |  | 0.6 |
| 0 | 0 (0%) | 0 (0%) |  |
| 1 | 18 (95%) | 16 (89%) |  |
| 2 | 1 (5.3%) | 2 (11%) |  |
| 3 | 0 (0%) | 0 (0%) |  |
| missing | 0 | 2 |  |
| Use of sleep medication |  |  | >0.9 |
| 0 | 18 (95%) | 18 (100%) |  |
| 1 | 1 (5.3%) | 0 (0%) |  |
| 2 | 0 (0%) | 0 (0%) |  |
| 3 | 0 (0%) | 0 (0%) |  |
| missing | 0 | 2 |  |
| Daytime dysfunction |  |  | 0.3 |
| 0 | 3 (16%) | 7 (39%) |  |
| 1 | 11 (58%) | 7 (39%) |  |
| 2 | 4 (21%) | 4 (22%) |  |
| 3 | 1 (5.3%) | 0 (0%) |  |
| missing | 0 | 2 |  |
| Global PSQI | 5.00 (1.00,11.00) | 3.50 (2.00,8.00) | 0.15 |
| missing | 0 | 2 |  |

<sup>1</sup> n (%); Median (Min,Max) <sup>2</sup> \*p<0.05; \*\*p<0.01; \*\*\*p<0.001

### Supplementary Figures

**Figure S1.**
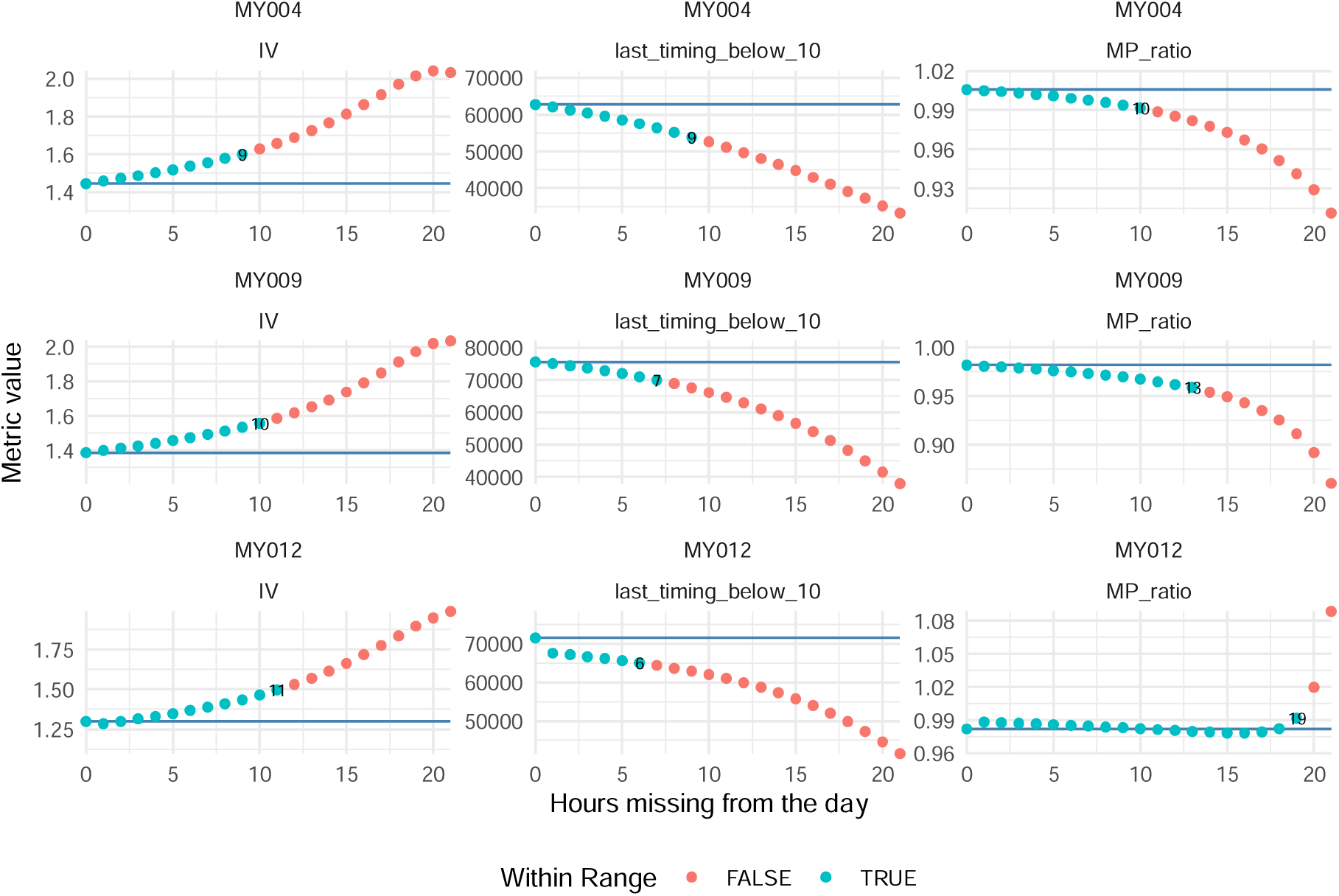
Summary of the bootstrapping procedure to determine an acceptable threshold of daily missing. Data based on three individual participants from the Malaysia site (MY004, MY009, MY012). Three variables were randomly chosen (IV, last time below 10 lux mel EDI, MP ratio). The blue horizontal line shows the average value of data across the full dataset, the blue rectangle a 95% confidence interval around that average. Dots show the average value across all bootstraps of that respective threshold value. A blue dot indicates that all 10^4^ bootstraps lie within the 95% confidence interval of the full dataset. A red dot indicates that at least one bootstrap lies outside of the 95% confidence interval. Abbreviations: IV; Interdaily variability; MP ratio, Melanopic/photopic ratio.

**Figure S2.**
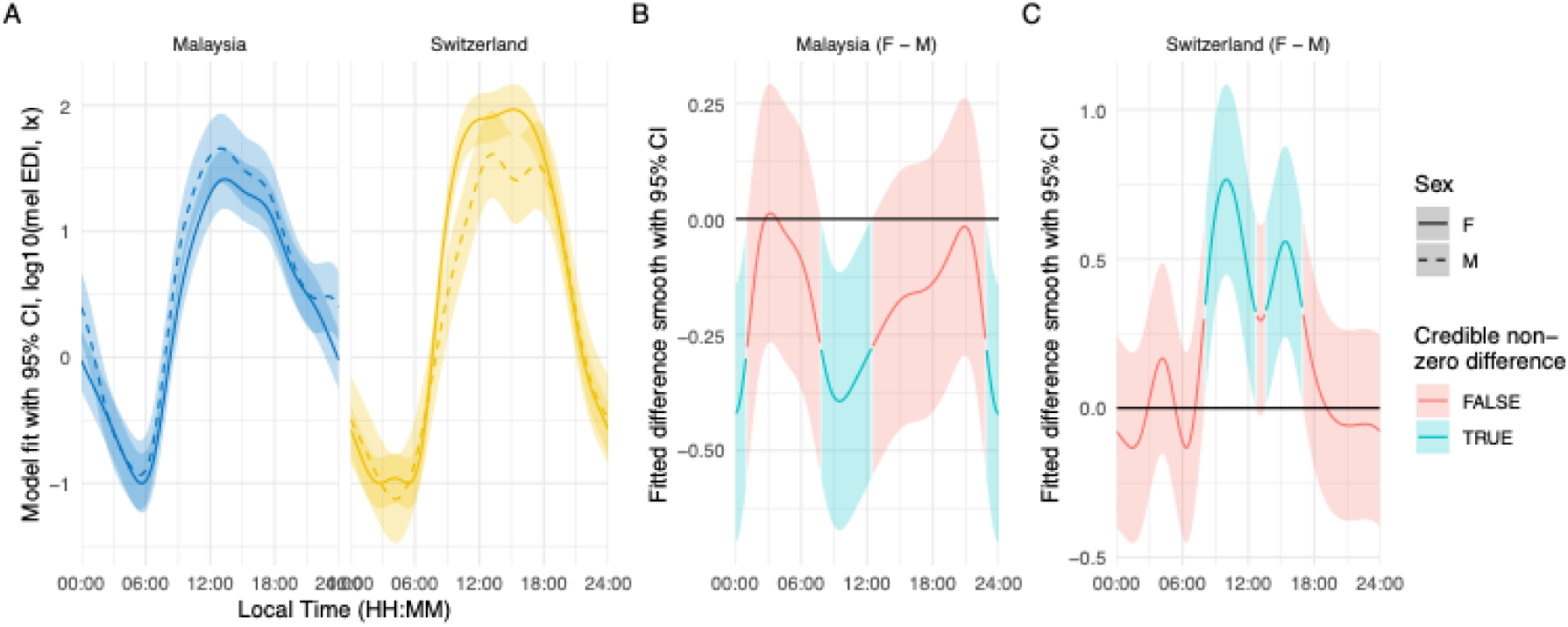
Sex differences in light exposure patterns. Results from a generalized additive mixed model looking at light exposure patterns across the day, dependent on site and sex. A: Model predictions for log10 melanopic EDI depending on local time, for both sites (panels) and sexes (solid and dashed lines). Bands indicate 95% confidence intervals. B: Difference smooth of log10 melanopic EDI between females and males in Malaysia. Red lines and bands indicate a non-significant differences, blue lines and bands a significant one. Values above 0 indicate higher illuminance for females at a given time, values below 0 vice versa. C: like B, but for Switzerland.

**Figure S3.**
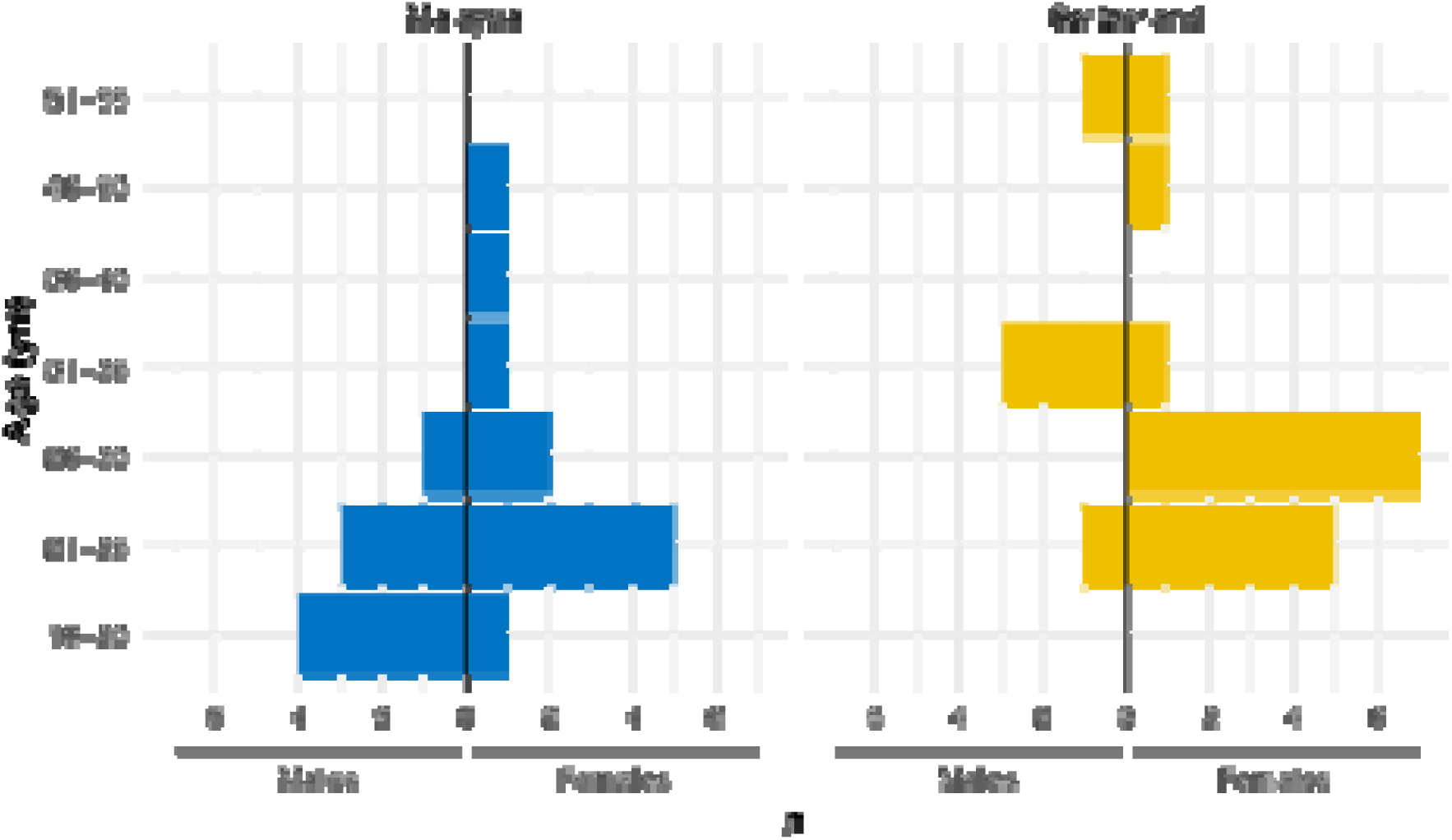
Distribution of age and sex between the sites. 2-way histograms for both Malaysia and Switzerland sites, showing the distribution of age in 5-year bins (+18 to 20yrs). Left of each central vertical line, male counts are shown, and females are on the right.

**Figure S4.**
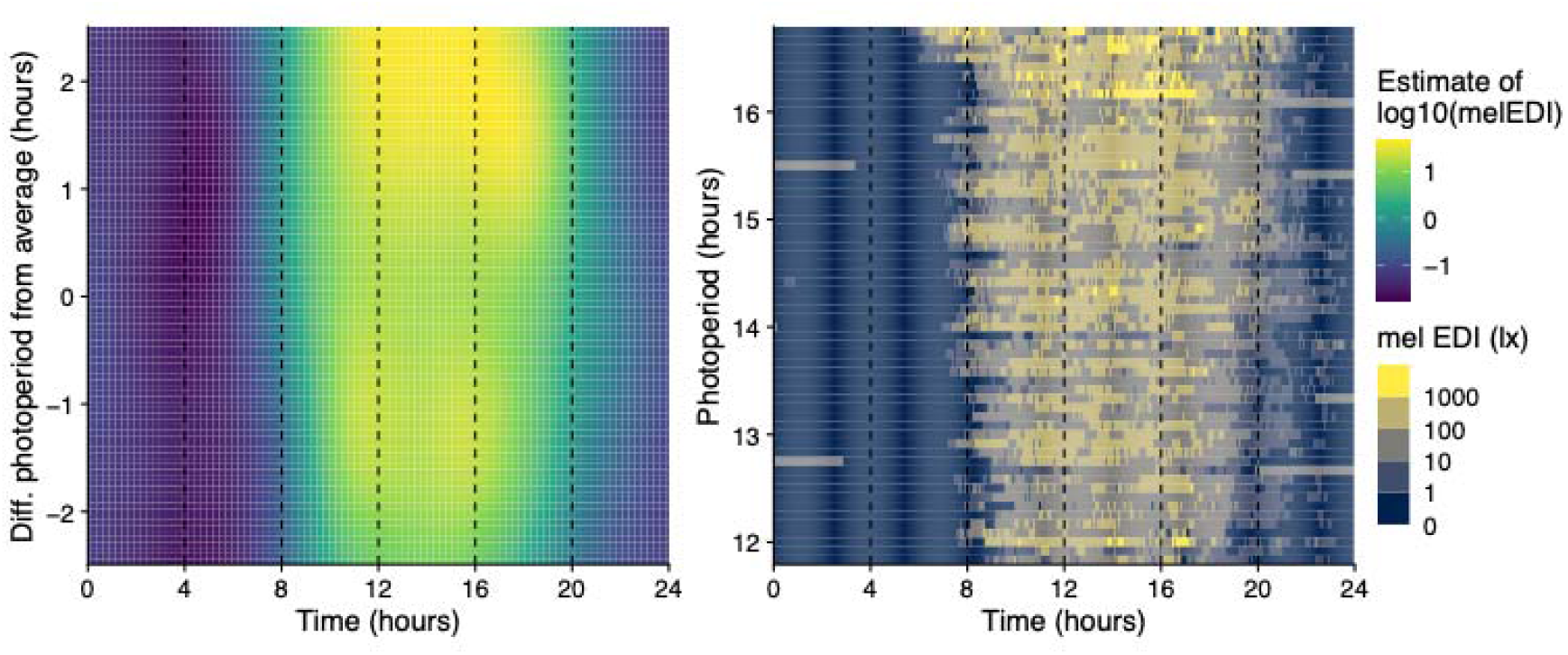
Light exposure patterns depending on photoperiod duration. Left: Model predictions from a generalised additive mixed-effect model of photoperiod duration, plotted as a difference in hours from the average, against the local time (in hours). Colour indicates log10 melanopic EDI. Right: Same as A, but based on measurement values, and absolute photoperiod duration. Measurements are binned into colour steps of log10 melanopic EDI.

**Figure S5.**
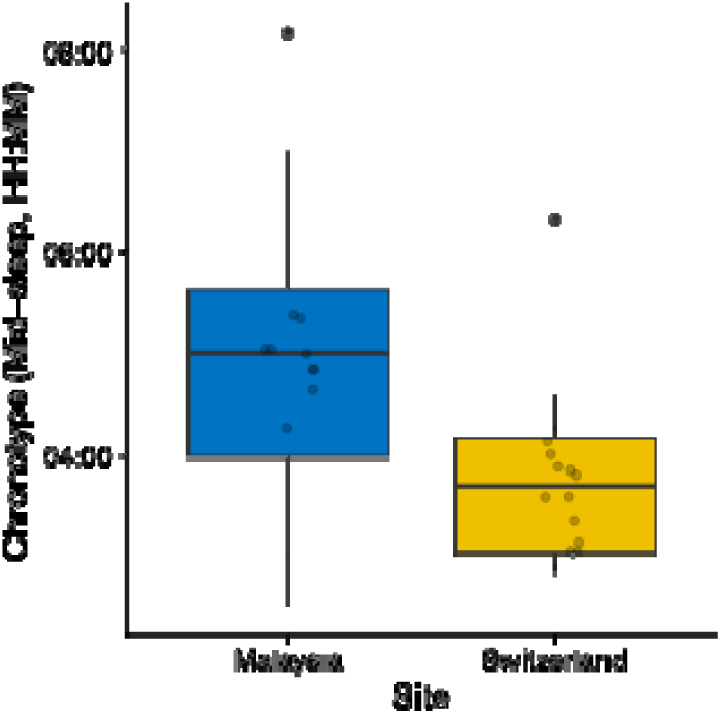
Chronotype distribution across sites. Boxplot with indicated individual values of chronotype by site. Chronotype is provided as the mid-sleep timing (HH:MM). Mid-sleep was 4:57 a.m. ±1.4 hours in Malaysia, and 3:45 a.m. ±0.8 hours in Switzerland. The difference is significant (*p = 0.003*).

**Figure S6.**
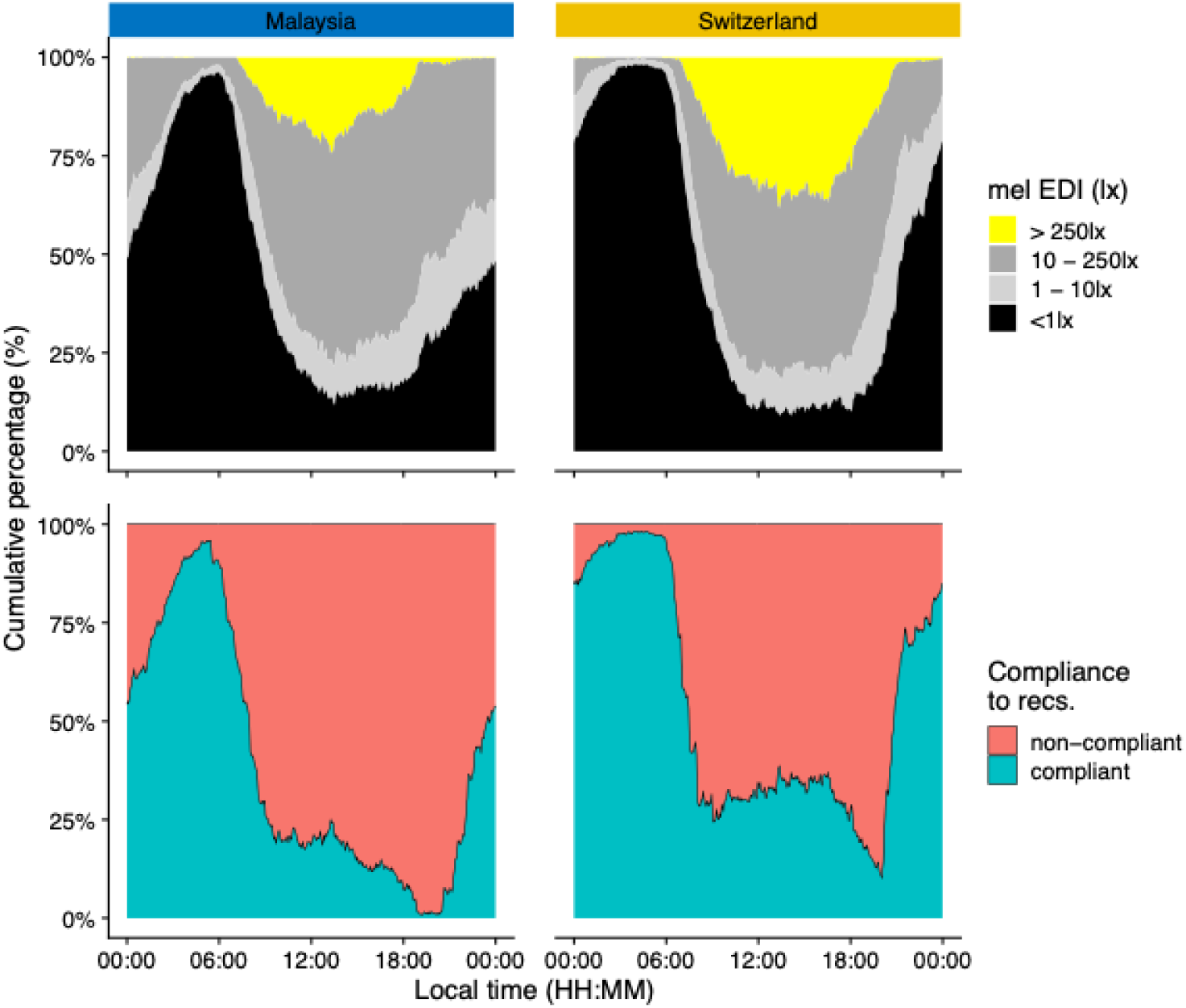
Light exposure in relation to the Brown recommendations for healthy lighting. Top panels: cumulative relative distribution of light exposure to recommended levels across the day; up to 1 lx mel EDI for sleep, up to 10 lx mel EDI for the evening, and above 250 lx for daytime. Bottom panels: Compliance to the recommendations across the day across all participants.

## Full Analysis Document

The full analysis document can be found on our GitHub page: https://tscnlab.github.io/BillerEtAl_JExpoSciEnvironEpidemiol_2025/

